## Supplementary Material 1 for "Synergistic genomic mechanisms of adaptation to ocean acidification in a coral holobiont"

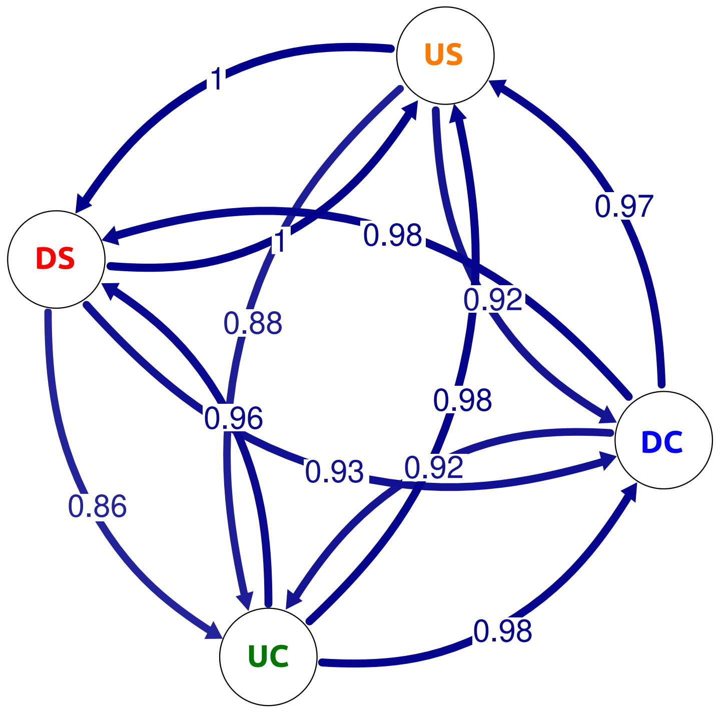


**Supplementary Material 1:** $G_{ST}$ Migration network. Circles represent sampling sites: red DS: Dobu Seep; blue DC, Dobu Control; orange US, Upa-Upasina Seep; green UC, Upa-Upasina Control.
