## Supplementary Material 2 for "Synergistic genomic mechanisms of adaptation to ocean acidification in a coral holobiont"

**Supplementary Material 2. Complete list of candidate adaptive SNPs with their positions, names in the A. millepora reference genome, annotations, SnpEff results and E-value of the blast searches for the SNPs that fell in intergenic regions**

| Chromosome | Position | Name of the gene | Annotation | SnpEff annotation | E-value of the blast searches |
|---|---|---|---|---|---|
| chr1 | 568394 | Amillepora00044 | Twitchin | synonymous | |
| chr1 | 1038464 | Amillepora00093 | Calreticulin | 3-prime-UTR | |
| chr1 | 1044515 | Amillepora00097 | Protein disabled | 5-prime-UTR | |
| chr1 | 1044530 | Amillepora00097 | Protein disabled | 5_prime_UTR_premature_start_codon_gain_variant | |
| chr1 | 1683848 | Intergenic region | Acropora millepora ribosomal RNA large subunit methyltransferase E-like | | 5.00E-114 |
| chr1 | 2914154 | Amillepora00278 | Actin-related protein 5 | synonymous | |
| chr1 | 2914167 | Amillepora00278 | Actin-related protein 5 | synonymous | |
| chr1 | 5053469 | Intergenic region | Acropora millepora negative elongation factor E-like (LOC114950754), mRNA | | 0 |
| chr1 | 5329212 | Intergenic region | Acropora millepora Na(+)/H(+) exchanger beta-like | | 3.00E-104 |
| chr1 | 7386545 | Amillepora00666 | Rootletin | Exon, AA change: Ser515Gly | |
| chr1 | 8649199 | Intergenic region | Acropora millepora probable E3 ubiquitin-protein ligase makorin-1 | | 0 |
| chr1 | 8752443 | Amillepora00758 | Complement C3 | Intron | |
| chr1 | 8755148 | Amillepora00758 | Complement C3 | synonymous | |
| chr1 | 9611781 | Intergenic region | Acropora millepora dynein axonemal assembly factor 5-like (LOC114962884), mRNA | | 0 |
| chr1 | 11207173 | Intergenic region | --- | | |
| chr1 | 11207280 | Intergenic region | --- | | |
| chr1 | 11621523 | Intergenic region | Acropora millepora charged multivesicular body protein 3-like (LOC114967980), mRNA | | 6.00E-173 |
| chr1 | 12924016 | Intergenic region | Acropora digitifera uncharacterized LOC107343730 (LOC107343730), transcript variant X5, mRNA | | 0 |
| chr1 | 13998242 | Intergenic region | DNA repair protein complementing XP-A cells homolog | 2 bases Upstream | |
| chr1 | 14719608 | Amillepora01328 | Valine--tRNA ligase | Exon synonymous variant | |
| chr1 | 16612927 | Amillepora01474 | Acropora millepora tumor necrosis factor receptor superfamily member 16-like | 3-prime-UTR | 0 |
| chr1 | 16719005 | Intergenic region | Acropora millepora rab GTPase-activating protein 1-like | | |
| chr1 | 16998322 | Amillepora01509 | Tachylectin-2 | Exon, AA change: Ser264Gly | |
| chr1 | 16998347 | Intergenic region | Tachylectin-2 | | |
| chr1 | 16998363 | Intergenic region | Tachylectin-2 | | |
| chr1 | 17957256 | Amillepora01580 | Zinc finger CCCH-type antiviral protein 1 | 3-prime-UTR | |
| chr1 | 17958867 | Intergenic region | Acropora digitifera poly [ADP-ribose] polymerase 12-like | | 0 |
| chr1 | 18060546 | Amillepora01593 | Acropora millepora uncharacterized LOC114955001 | Intron | 0 |
| chr1 | 19617261 | Intergenic region | Acropora millepora fructose-1,6-bisphosphatase 1-like | | 3.00E-19 |
| chr1 | 19951791 | Amillepora01782 | Stomatin-like protein 2, mitochondrial | 3-prime-UTR | |
| chr1 | 20063107 | Intergenic region | Acropora millepora ADP-ribosylation factor-like protein 15 | | 3.00E-157 |
| chr1 | 20843187 | Intergenic region | Acropora millepora spatacsin-like | | 0 |
| chr1 | 30732002 | Intergenic region | Acropora millepora peptidyl alpha-hydroxylating monooxygenase (PHM) | | 0 |
| chr1 | 32325594 | Amillepora02704 | COUP transcription factor 2 | 3-prime-UTR | |
| chr1 | 32827163 | Intergenic region | Acropora digitifera transmembrane protein 163-like | | 1.00E-130 |
| chr1 | 32828045 | Intergenic region | Acropora digitifera transmembrane protein 163-like | | 0 |
| chr1 | 33073572 | Intergenic region | Acropora millepora protocadherin Fat 4-like | | 0 |
| chr1 | 34103026 | Intergenic region | Acropora millepora Golgi phosphoprotein 3-like | | 0 |
| chr1 | 34148371 | Amillepora02847 | Ubiquitin carboxyl-terminal hydrolase 3 | Exon synonymous variant | |
| chr1 | 35264398 | Amillepora02941 | Thrombospondin-3 | Exon, AA change: Met277Ile | |
| chr1 | 35385011 | Amillepora02951 | Dynein intermediate chain 2, ciliary | 3-prime-UTR | |
| chr1 | 35800287 | Intergenic region | Acropora millepora uncharacterized LOC114973780 | | 0 |
| chr1 | 37497226 | Amillepora03106 | Centrosomal protein of 41 kDa | 3-prime-UTR | |
| chr1 | 38903260 | Intergenic region | Acropora millepora ninein-like | | 0 |
| chr1 | 38973443 | Intergenic region | Acropora digitifera midasin-like | | 0 |
| chr2 | 559818 | Intergenic region | Acropora millepora solute carrier family 40 member 1-like | | 7.00E-172 |
| chr2 | 1100299 | Intergenic region | Acropora digitifera casein kinase I isoform alpha-like | | 3.00E-61 |
| chr2 | 1525066 | Amillepora03361 | L-seryl-tRNA(Sec) kinase | Exon, AA change: Glu376Asp | |
| chr2 | 2961772 | Amillepora03480 | Heparanase | 3-prime-UTR | |
| chr2 | 3045560 | Amillepora03482 | *Golgi-associated plant pathogenesis-related protein 1* | synonymous | |
| chr2 | 3046213 | Amillepora03482 | *Golgi-associated plant pathogenesis-related protein 2* | synonymous | |
| chr2 | 3219420 | Amillepora03492 | CREB3 regulatory factor | 3-prime-UTR | |
| chr2 | 3761552 | Amillepora03530 | Dipeptidyl peptidase 1 | Intron | |
| chr2 | 3966029 | Amillepora03553 | Structural maintenance of chromosomes protein 3 | synonymous | |
| chr2 | 3992964 | Amillepora03556 | Uncharacterized WD repeat-containing protein alr2800 | 3-prime-UTR | |
| chr2 | 4762356 | Amillepora03625 | Glycosylphosphatidylinositol anchor attachment 1 protein | 3-prime-UTR | |
| chr2 | 4777662 | Intergenic region | Acropora millepora cytochrome c1, heme protein, mitochondrial-like | | 2.00E-114 |
| chr2 | 5066179 | Intergenic region | Acropora millepora CDK5 regulatory subunit-associated protein 2-like | | 9.00E-135 |
| chr2 | 5435429 | Intergenic region | Acropora millepora hexokinase-2-like | | 0 |
| chr2 | 5610872 | Intergenic region | Acropora millepora xaa-Pro aminopeptidase 1-like | | 0 |
| chr2 | 5811932 | Amillepora03714 | Catenin alpha-2 | 3-prime-UTR | |
| chr2 | 5969212 | Amillepora03732 | Probable prolyl 4-hydroxylase 7 | 3-prime-UTR | |
| chr2 | 6624582 | Intergenic region | --- | | |
| chr2 | 6624614 | Intergenic region | --- | | |
| chr2 | 9005067 | Amillepora03938 | Neuferricin | 3-prime-UTR | |
| chr2 | 15018462 | Amillepora04338 | -- | 5-prime-UTR | |
| chr2 | 15111325 | Amillepora04345 | Elongation factor 1-gamma | synonymous | |
| chr2 | 15332951 | Amillepora04364 | Cilia- and flagella-associated protein 36 | synonymous | |
| chr2 | 15433795 | Amillepora04376 | Tenascin-X | 3-prime-UTR | |
| chr2 | 15584711 | Intergenic region | Acropora digitifera uncharacterized LOC107345063 | | 0 |
| chr2 | 15584826 | Intergenic region | Acropora digitifera uncharacterized LOC107345063 | | 0 |
| chr2 | 15584870 | Intergenic region | Acropora digitifera uncharacterized LOC107345063 | | 0 |
| chr2 | 16375397 | Intergenic region | Acropora millepora 60S ribosomal protein L44-like | | 3.00E-51 |
| chr2 | 16375432 | Intergenic region | Acropora millepora 60S ribosomal protein L44-like | | 3.00E-51 |
| chr2 | 16676923 | Intergenic region | Acropora millepora transient receptor potential cation channel subfamily M member-like 2 | | 0 |
| chr2 | 17338566 | Amillepora04500 | WD40 repeat-containing protein SMU1 | 3-prime-UTR | |
| chr2 | 17338600 | Amillepora04500 | WD40 repeat-containing protein SMU1 | 3-prime-UTR | |
| chr2 | 18783694 | Amillepora04617 | Coiled-coil domain-containing protein 149 | 3-prime-UTR | |
| chr2 | 19181417 | Amillepora04653 | Acropora millepora uncharacterized LOC114965138 | 3-prime-UTR | 0 |
| chr2 | 19206162 | Amillepora04654 | EF-hand calcium-binding domain-containing protein 6 | synonymous | |
| chr2 | 19583718 | Amillepora04680 | Mitochondrial uncoupling protein 2 | 3-prime-UTR | |
| chr2 | 19958850 | Amillepora04702 | myosin-2 heavy chain, non muscle-like isoform X2 [Stylophora pistillata] | synonymous | 2.00E-31 |
| chr2 | 19958881 | Amillepora04702 | myosin-2 heavy chain, non muscle-like isoform X2 [Stylophora pistillata] | 3-prime-UTR | 2.00E-31 |
| chr2 | 19993016 | Intergenic region | Acropora millepora lysosomal acid lipase/cholesteryl ester hydrolase-like | | 0 |
| chr2 | 20016051 | Intergenic region | Acropora millepora serine/threonine-protein phosphatase 6 regulatory subunit 3-like | | 0 |
| chr2 | 20016949 | Intergenic region | Acropora millepora serine/threonine-protein phosphatase 6 regulatory subunit 3-like | | 0 |
| chr2 | 20017144 | Amillepora04708 | Pantetheinase | synonymous | |
| chr2 | 20391253 | Intergenic region | Acropora digitifera adhesion G-protein coupled receptor D1-like | | 0 |
| chr2 | 20821826 | Amillepora04771 | Acropora millepora uncharacterized LOC114965354 | Exon, AA change: Thr390Met | |
| chr2 | 21154302 | Intergenic region | Acropora millepora dual specificity mitogen-activated protein kinase kinase 5-like | | 2.00E-97 |
| chr2 | 21514317 | Intergenic region | Acropora millepora organic cation transporter protein-like | | 0 |
| chr2 | 21929156 | Amillepora04880 | Acropora millepora golgin subfamily B member 1-like | Exon, AA change: Glu1679Gln | 0 |
| chr2 | 21929864 | Amillepora04880 | Acropora millepora golgin subfamily B member 1-like | Exon, AA change: Ile1915Val | 0 |
| chr2 | 22074518 | Amillepora04880 | Acropora millepora golgin subfamily B member 1-like | Exon, AA change: Trp209Leu | 0 |
| chr2 | 22463916 | Amillepora04913 | Protein lin-54 homolog | 3-prime-UTR | |
| chr2 | 22584528 | Amillepora04922 | Carbonic anhydrase 2 | Intron | |
| chr2 | 22793310 | Amillepora04935 | Sodium- and chloride-dependent GABA transporter 1 | Intron | |
| chr2 | 23057113 | Amillepora04958 | Major facilitator superfamily domain-containing protein 6 | 3-prime-UTR | |
| chr2 | 24101619 | Amillepora05037 | Cyclin-dependent kinase inhibitor 1B | 3-prime-UTR | |
| chr2 | 24186313 | Intergenic region | --- | | |
| chr2 | 24246921 | Amillepora05049 | Synaptotagmin-9 | Intron | |
| chr2 | 24864042 | Intergenic region | Acropora millepora uncharacterized LOC114967882 | | 0 |
| chr2 | 26138478 | Intergenic region | Acropora millepora guanine nucleotide-binding protein G(q) subunit alpha-like | | 0 |
| chr2 | 26532719 | Amillepora05264 | Coiled-coil domain-containing protein 186 | Intron | |
| chr2 | 26533131 | Amillepora05264 | Coiled-coil domain-containing protein 186 | Intron | |
| chr2 | 26533155 | Amillepora05264 | Coiled-coil domain-containing protein 186 | Intron | |
| chr2 | 27746840 | Amillepora05349 | 5-oxoprolinase | 3-prime-UTR | |
| chr2 | 27869424 | Amillepora05366 | NADH dehydrogenase [ubiquinone] 1 beta subcomplex subunit 9 | 3-prime-UTR | |
| chr2 | 28665845 | Intergenic region | --- | | |
| chr2 | 29349391 | Intergenic region | --- | | |
| chr2 | 31055981 | Amillepora05615 | Beta-1,4-galactosyltransferase 1 | synonymous | |
| chr2 | 31441612 | Amillepora05646 | Acropora millepora tetratricopeptide repeat protein 28-like | 3-prime-UTR | 0 |
| chr2 | 32134310 | Amillepora05687 | Hematopoietically-expressed homeobox protein hhex | 3-prime-UTR | |
| chr2 | 32354512 | Amillepora05711 | Dehydrogenase/reductase SDR family member 7 | 3-prime-UTR | |
| chr2 | 32456006 | Amillepora05723 | Dynactin subunit 1 | 3-prime-UTR | |
| chr2 | 32456030 | Amillepora05723 | Dynactin subunit 1 | 3-prime-UTR | |
| chr2 | 33292425 | Amillepora05797 | 9,11-endoperoxide prostaglandin H2 reductase | 3-prime-UTR | |
| chr2 | 33297233 | Amillepora05797 | 9,11-endoperoxide prostaglandin H2 reductase | synonymous | |
| chr2 | 34311132 | Intergenic region | Acropora millepora uncharacterized LOC114967523 | | 4.00E-59 |
| chr2 | 34778321 | Amillepora05919 | Tyrosine-protein phosphatase non-receptor type 13 | Intron | |
| chr2 | 34909495 | Amillepora05936 | Tight junction protein ZO-2 | Intron | |
| chr2 | 35236874 | Intergenic region | Acropora digitifera uncharacterized LOC107340049 | | 2.00E-93 |
| chr2 | 36315950 | Intergenic region | --- | | |
| chr2 | 36317233 | Intergenic region | Acropora digitifera protein N-lysine methyltransferase METTL21A-like | | 9.00E-48 |
| chr2 | 36368288 | Amillepora06063 | Coiled-coil domain-containing protein 180 | 3-prime-UTR | |
| chr2 | 36856167 | Amillepora06104 | Calumenin-A | 3-prime-UTR | |
| chr2 | 37003630 | Intergenic region | Acropora millepora torsin-1A-like | | 0 |
| chr2 | 37004245 | Intergenic region | Acropora millepora ubiquitin carboxyl-terminal hydrolase 20-like | | 0 |

| | | | | | |
|---|---|---|---|---|---|
| chr3 | 170761 | Amillepora21892 | Acropora millepora uncharacterized LOC114966735 | 3-prime-UTR | 0 |
| chr3 | 170890 | Amillepora21892 | Acropora millepora uncharacterized LOC114966735 | 3-prime-UTR | 0 |
| chr3 | 276790 | Amillepora21901 | Acropora millepora bromodomain adjacent to zinc finger domain protein 2B-like | 3-prime-UTR | 5.00E-49 |
| chr3 | 941364 | Amillepora21960 | Replication protein A 70 kDa DNA-binding subunit | 3-prime-UTR | |
| chr3 | 959737 | Amillepora21966 | Very-long-chain enoyl-CoA reductase | 3-prime-UTR | |
| chr3 | 959741 | Amillepora21966 | Very-long-chain enoyl-CoA reductase | 3-prime-UTR | |
| chr3 | 972454 | Intergenic region | Acropora millepora tetratricopeptide repeat protein 41-like | | 0 |
| chr3 | 1230378 | Amillepora21990 | Protein phosphatase 1 regulatory subunit 42 | 3-prime-UTR | |
| chr3 | 1603748 | Amillepora22025 | Ubiquitin recognition factor in ER-associated degradation protein | 3-prime-UTR | |
| chr3 | 1694437 | Amillepora22034 | Acropora millepora serine/arginine repetitive matrix protein 2-like | 3-prime-UTR | 0 |
| chr3 | 1694447 | Amillepora22034 | Acropora millepora serine/arginine repetitive matrix protein 2-like | 3-prime-UTR | 0 |
| chr3 | 1853631 | Amillepora22048 | Ovarian abundant message protein | Exon, AA change: Asn638Ser | |
| chr3 | 2105604 | Amillepora22078 | Disks large-associated protein 5 | 3-prime-UTR | |
| chr3 | 2105772 | Amillepora22078 | Disks large-associated protein 5 | 3-prime-UTR | |
| chr3 | 2106025 | Amillepora22078 | Disks large-associated protein 5 | 3-prime-UTR | |
| chr3 | 2106049 | Amillepora22078 | Disks large-associated protein 5 | 3-prime-UTR | |
| chr3 | 2106102 | Amillepora22078 | Disks large-associated protein 5 | 3-prime-UTR | |
| chr3 | 2106492 | Amillepora22078 | Disks large-associated protein 5 | 3-prime-UTR | |
| chr3 | 2163776 | Amillepora22082 | Probable ATP-dependent RNA helicase DDX43 | 3-prime-UTR | |
| chr3 | 2169955 | Amillepora22084 | Tropomyosin-2 | Exon, AA change: Lys274Glu | |
| chr3 | 2703390 | Intergenic region | Acropora digitifera protein-L-isoaspartate(D-aspartate) O-methyltransferase-like | | 0 |
| chr3 | 2705842 | Intergenic region | Acropora millepora protein-L-isoaspartate(D-aspartate) O-methyltransferase-like | | 0 |
| chr3 | 2759031 | Intergenic region | Acropora millepora septin-2B-like | | |
| chr3 | 2760331 | Amillepora22151 | Septin-2 | 3-prime-UTR | |
| chr3 | 2783766 | Intergenic region | Acropora digitifera uncharacterized LOC107341289 | | 0 |
| chr3 | 2876473 | Amillepora22163 | Max-binding protein MNT | 3-prime-UTR | |
| chr3 | 2902332 | Intergenic region | Acropora digitifera probable E3 ubiquitin-protein ligase HECTD4 | | 4.00E-69 |
| chr3 | 3561079 | Intergenic region | --- | | |
| chr3 | 4228078 | Amillepora22273 | Ran-specific GTPase-activating protein | Exon synonymous variant | |
| chr3 | 4229120 | Amillepora22274 | Developmentally-regulated GTP-binding protein 1 | 3-prime-UTR | |
| chr3 | 4229227 | Amillepora22274 | Developmentally-regulated GTP-binding protein 1 | 3-prime-UTR | |
| chr3 | 4398295 | Amillepora22291 | P2X purinoceptor 4 | synonymous | |
| chr3 | 4631570 | Amillepora22318 | Acropora millepora golgin subfamily A member 3-like | 5_prime_UTR | 0 |
| chr3 | 4636289 | Amillepora22318 | – | Intron | |
| chr3 | 4636317 | Amillepora22318 | – | Intron | |
| chr3 | 4898052 | Intergenic region | Acropora millepora protein LIAT1-like | | 0 |
| chr3 | 5023205 | Amillepora22359 | Dystonin | 3-prime-UTR | |
| chr3 | 5041034 | Amillepora22360 | Plectin | 3-prime-UTR | |
| chr3 | 5089865 | Amillepora22360 | Plectin | Intron | |
| chr3 | 5094777 | Amillepora22360 | Plectin | synonymous | |
| chr3 | 5132596 | Intergenic region | --- | | |
| chr3 | 5398555 | Intergenic region | Acropora millepora G patch domain-containing protein 3-like | | 0 |
| chr3 | 5459484 | Amillepora22378 | Lysine-specific demethylase 2B | Intron | |
| chr3 | 5937459 | Amillepora22405 | Protein phosphatase 1E | 3-prime-UTR | |
| chr3 | 7004689 | Amillepora22515 | Peripheral-type benzodiazepine receptor-associated protein 1 | synonymous | |
| chr3 | 8010758 | Intergenic region | Acropora digitifera uncharacterized LOC107356740 | | 4.00E-110 |
| chr3 | 8909289 | Intergenic region | Acropora millepora collagen alpha-2(IX) chain-like | | 4.00E-70 |
| chr3 | 8954774 | Amillepora22668 | Aquaporin AQPAn.G | 3-prime-UTR | |
| chr3 | 8954965 | Amillepora22668 | Aquaporin AQPAn.G | 3-prime-UTR | |
| chr3 | 14819058 | Amillepora23090 | Poly(U)-specific endoribonuclease-B | 3-prime-UTR | |
| chr3 | 15948124 | Amillepora23165 | Uncharacterized skeletal organic matrix protein 6 | Intron | |
| chr3 | 16061855 | Intergenic region | Acropora millepora protein ANKUB1-like | | 0 |
| chr3 | 16475553 | Intergenic region | Acropora millepora uncharacterized LOC114955923 | | 0 |
| chr3 | 16750458 | Intergenic region | Acropora digitifera uncharacterized LOC107348935 | | 0 |
| chr3 | 17249838 | Amillepora23272 | Proton-coupled amino acid transporter 1 | 3-prime-UTR | |
| chr3 | 17277143 | Amillepora23278 | Actin, cytoplasmic | 3-prime-UTR | |
| chr3 | 17301480 | Amillepora23283 | Acropora digitifera uncharacterized LOC107346909 | synonymous | 2.00E-152 |
| chr3 | 18211715 | Intergenic region | --- | | |
| chr3 | 18336843 | Amillepora23368 | – | Exon, AA change: Lys111Arg | |
| chr3 | 18708862 | Amillepora23389 | Acropora digitifera uncharacterized LOC107335365 | 5_prime_UTR | 4.00E-156 |
| chr3 | 18795684 | Amillepora23397 | Nucleoprotein TPR | synonymous | |
| chr3 | 19355004 | Amillepora23441 | Acropora millepora uncharacterized LOC114958002 | Exon, AA change: Glu41Asp | 1.00E-42 |
| chr3 | 19422635 | Amillepora23448 | Protein DEK | Exon-Intron splice acceptor variant | |
| chr4 | 362817 | Intergenic region | Acropora millepora GTP-binding protein Di-Ras1-like | | 0 |
| chr4 | 721923 | Amillepora08644 | Piezo-type mechanosensitive ion channel component 2 | 3-prime-UTR | |
| chr4 | 1708973 | Intergenic region | Acropora millepora serine/threonine-protein kinase 35-like | | 0 |
| chr4 | 3265784 | Amillepora08867 | Chromosome transmission fidelity protein 8 homolog | 3-prime-UTR | |
| chr4 | 3295940 | Amillepora08869 | GATOR complex protein WDR59 | 3-prime-UTR | |
| chr4 | 5235614 | Intergenic region | Acropora millepora probable D-lactate dehydrogenase, mitochondrial | | 0 |
| chr4 | 6273822 | Intergenic region | Acropora millepora galanin receptor type 1-like | | 0 |
| chr4 | 9411379 | Intergenic region | Acropora millepora tetratricopeptide repeat protein 28-like | | 3.00E-147 |
| chr4 | 11009882 | Amillepora09499 | Microfibrillar-associated protein 1 | 3-prime-UTR | |
| chr4 | 11009948 | Amillepora09499 | Microfibrillar-associated protein 1 | 3-prime-UTR | |
| chr4 | 11014906 | Amillepora09499 | Microfibrillar-associated protein 1 | Exon, AA change: Arg232Gln | |
| chr4 | 11294356 | Intergenic region | Ubiquitin carboxyl-terminal hydrolase MINDY-1 | | |
| chr4 | 11848347 | Amillepora09566 | Acetolactate synthase-like protein | 3-prime-UTR | |
| chr4 | 12682963 | Amillepora09624 | Dipeptidase 1 | synonymous | |
| chr4 | 12683010 | Amillepora09624 | Dipeptidase 1 | 3-prime-UTR | |
| chr4 | 13514058 | Amillepora09679 | 60S ribosomal protein L12 | 3-prime-UTR | |
| chr4 | 13901015 | Amillepora09700 | Telomerase protein component 1 | synonymous | |
| chr4 | 13977790 | Amillepora09708 | Sorting nexin-33 | synonymous | |
| chr4 | 14913025 | Intergenic region | Acropora millepora tyrosine-protein kinase Fer-like | | 5.00E-180 |
| chr4 | 15259153 | Amillepora09817 | Epidermal growth factor receptor kinase substrate 8-like protein 1 | 3-prime-UTR | |
| chr4 | 15289226 | Intergenic region | Acropora millepora plastin-3-like | | 0 |
| chr4 | 16749975 | Amillepora09932 | Alpha-actinin-1 | synonymous | |
| chr4 | 17301308 | Amillepora09972 | WD repeat-containing protein 76 | 3-prime-UTR | |
| chr4 | 17594372 | Intergenic region | Acropora millepora uncharacterized LOC114967123 | | 0 |
| chr4 | 18157729 | Amillepora10024 | Zinc finger CCCH domain-containing protein 8 | synonymous | |
| chr4 | 18218514 | Amillepora10033 | Splicing factor 3A subunit 1 | synonymous | |
| chr4 | 18842659 | Amillepora10089 | Mitogen-activated protein kinase kinase kinase 19 | Exon, AA change: Gly942Ser | |
| chr4 | 18849332 | Amillepora10089 | Mitogen-activated protein kinase kinase kinase 19 | 3-prime-UTR | |
| chr4 | 18987332 | Amillepora10102 | Transcription factor HES-1 | Intron | |
| chr4 | 18987564 | Amillepora10102 | Transcription factor HES-1 | Intron | |
| chr4 | 18996141 | Amillepora10103 | Gamma-aminobutyric acid type B receptor subunit 2 | 3-prime-UTR | |
| chr4 | 19822899 | Intergenic region | Acropora millepora intracellular protein transport protein USO1-like | | 0 |
| chr4 | 19978390 | Intergenic region | --- | | |
| chr4 | 20427311 | Amillepora10216 | Acropora millepora TNF receptor-associated factor 6-like | Intron | 0 |
| chr4 | 20885024 | Intergenic region | Acropora millepora potassium voltage-gated channel subfamily A member 7-like | | 0 |
| chr4 | 21519233 | Amillepora10346 | Probable ATP-dependent RNA helicase DDX31 | 5_prime_UTR_premature_start_codon_gain_variant | |
| chr4 | 22148290 | Intergenic region | Acropora millepora cyclin-C-like | | 3.00E-179 |
| chr4 | 22296748 | Intergenic region | Acropora digitifera RNA-binding protein 24-like | | 3.00E-12 |
| chr4 | 22402126 | Intergenic region | Acropora digitifera omega-6 fatty acid desaturase, endoplasmic reticulum isozyme 1-like | | 0 |
| chr4 | 22524749 | Amillepora10420 | Solute carrier family 22 member 5 | 3-prime-UTR | |
| chr4 | 22703862 | Intergenic region | --- | | |
| chr4 | 23532575 | Amillepora10499 | Phosphoenolpyruvate carboxykinase | 3-prime-UTR | |
| chr4 | 24772360 | Intergenic region | Acropora digitifera post-GPI attachment to proteins factor 2-like | | 6.00E-26 |
| chr4 | 24966058 | Amillepora10588 | Splicing factor 3B subunit 3 | Intron | |
| chr4 | 25823323 | Intergenic region | Acropora millepora 40S ribosomal protein S14 | | 1.00E-56 |
| chr4 | 26720901 | Amillepora10714 | Transcription factor E2F5 | 3-prime-UTR | |
| chr4 | 26785517 | Intergenic region | Acropora millepora importin-9-like | | 4.00E-13 |
| chr4 | 27860664 | Amillepora10796 | GMP reductase 2 | 3-prime-UTR | |
| chr4 | 27860686 | Amillepora10796 | GMP reductase 2 | 3-prime-UTR | |
| chr4 | 27860701 | Amillepora10796 | GMP reductase 2 | 3-prime-UTR | |
| chr4 | 29290257 | Amillepora10913 | LIM domain and actin-binding protein 1 | synonymous | |
| chr5 | 235421 | Amillepora06175 | Epsin-2 | 3-prime-UTR | |
| chr5 | 785756 | Intergenic region | Acropora digitifera homeobox protein Nkx-2.5-like | | 0 |
| chr5 | 2000969 | Amillepora06348 | Leucine-rich repeat-containing protein 59 | 5_prime_UTR | |
| chr5 | 2643775 | Amillepora06397 | ER membrane protein complex subunit 3 | 3-prime-UTR | |
| chr5 | 2874304 | Amillepora06422 | SH3 domain-containing protein 19 | synonymous | |
| chr5 | 3211346 | Amillepora06449 | Dolichyl-diphosphooligosaccharide–protein glycosyltransferase subunit 1 | 3-prime-UTR | |
| chr5 | 3713925 | Amillepora06489 | Fibropellin-1 | Exon, AA change: Val1578Leu | |
| chr5 | 4023020 | Intergenic region | Acropora millepora uncharacterized LOC114946839 | | 8.00E-73 |
| chr5 | 4137975 | Intergenic region | Acropora millepora aldehyde dehydrogenase, mitochondrial-like | | 0 |
| chr5 | 4684860 | Intergenic region | Acropora millepora uncharacterized LOC114975098 | | 0 |
| chr5 | 4817463 | Amillepora06585 | E3 ubiquitin-protein ligase HERC2 | synonymous | |
| chr5 | 6313549 | Amillepora06732 | Protein FAM114A2 | Exon, AA change: Thr196Lys | |
| chr5 | 6332419 | Intergenic region | Acropora millepora eukaryotic translation initiation factor 5B-like | | 0 |
| chr5 | 6332449 | Intergenic region | Acropora millepora eukaryotic translation initiation factor 5B-like | | 2.00E-86 |
| chr5 | 7187705 | Amillepora06789 | Transmembrane protein 33 | 3-prime-UTR | |
| chr5 | 7627336 | Intergenic region | Acropora millepora tenascin-like | | 4.00E-37 |
| chr5 | 10585100 | Intergenic region | Acropora millepora calnexin-like | | 0 |

| | | | | | |
|---|---|---|---|---|---|
| chr5 | 10599624 | Amillepora07010 | Calnexin | Exon, AA change: Val572Gly | |
| chr5 | 12178760 | Intergenic region | --- | | |
| chr5 | 12178780 | Intergenic region | --- | | |
| chr5 | 13136729 | Intergenic region | Acropora digitifera DNA-directed RNA polymerase I subunit RPA49-like | | 9.00E-57 |
| chr5 | 15023464 | Amillepora07303 | Grainyhead-like protein 2 homolog | 3-prime-UTR | |
| chr5 | 15152180 | Amillepora07314 | Agrin | Exon, AA change: Pro183Ser | |
| chr5 | 15484788 | Intergenic region | Orbicella faveolata uncharacterized LOC110068546 | | 4.00E-32 |
| chr5 | 15616866 | Amillepora07352 | Acropora millepora uncharacterized LOC114961008 | synonymous | 3.00E-57 |
| chr5 | 18882073 | Amillepora07598 | V-type proton ATPase subunit S1 | synonymous | |
| chr5 | 19948066 | Intergenic region | Acropora millepora phosphoribosylformylglycinamidine synthase-like | | 0 |
| chr5 | 19948147 | Intergenic region | Acropora millepora phosphoribosylformylglycinamidine synthase-like | | 0 |
| chr5 | 20519851 | Amillepora07718 | UDP-N-acetylglucosamine--peptide N-acetylglucosaminyltransferase 110 kDa subunit | 3-prime-UTR | |
| chr5 | 22833746 | Intergenic region | Acropora millepora protein Spindly-B-like | | 4.00E-114 |
| chr5 | 22853579 | Intergenic region | Acropora digitifera methionine adenosyltransferase 2 subunit beta-like | | 0 |
| chr5 | 24234243 | Amillepora08001 | Amyloid-beta A4 precursor protein-binding family B member 1 | Exon, AA change: Arg205Ser | |
| chr5 | 24236015 | Amillepora08001 | Amyloid-beta A4 precursor protein-binding family B member 1 | Exon, AA change: His718Arg | |
| chr5 | 24236160 | Amillepora08001 | Amyloid-beta A4 precursor protein-binding family B member 1 | Exon, AA change: Lys766Asn | |
| chr5 | 24595746 | Intergenic region | Acropora digitifera thioredoxin domain-containing protein 9-like | | 0 |
| chr5 | 24600259 | Intergenic region | Acropora millepora inositol 1,4,5-trisphosphate receptor-like | | 0 |
| chr5 | 24759153 | Intergenic region | Acropora millepora uncharacterized LOC114960886 | | 0 |
| chr5 | 24956399 | Amillepora08054 | 3-hydroxyacyl-CoA dehydrogenase type-2 | synonymous | |
| chr5 | 24956469 | Amillepora08054 | 3-hydroxyacyl-CoA dehydrogenase type-2 | Intron | |
| chr5 | 25036780 | Amillepora08060 | Acropora millepora uncharacterized LOC114960918 | 3-prime-UTR | 0 |
| chr5 | 25115878 | Amillepora08069 | Guanylate cyclase soluble subunit beta-1 | Intron | |
| chr5 | 25232636 | Amillepora08079 | Acropora millepora optineurin-like | synonymous | 0 |
| chr5 | 26444396 | Amillepora08202 | Acropora digitifera uncharacterized LOC107344986 | Exon, AA change: Met199Ile | 1.00E-121 |
| chr5 | 26908952 | Intergenic region | Acropora millepora beta-lactamase-like protein 2 | | 8.00E-08 |
| chr5 | 26909008 | Intergenic region | Acropora millepora beta-lactamase-like protein 2 | | |
| chr5 | 26909027 | Intergenic region | Acropora millepora beta-lactamase-like protein 2 | | |
| chr5 | 26909035 | Intergenic region | Acropora millepora beta-lactamase-like protein 2 | | 1.00E-61 |
| chr5 | 27031860 | Intergenic region | Acropora millepora BTB/POZ domain-containing protein KCTD8-like | | 0 |
| chr5 | 27463752 | Amillepora08287 | Pericentriolar material 1 protein | synonymous | |
| chr5 | 29320441 | Amillepora08401 | Protein Wnt-16 | 3-prime-UTR | |
| chr5 | 29352732 | Amillepora08404 | Acropora millepora uncharacterized LOC122959985 | 5_prime_UTR_premature_start_codon_gain_variant | 0 |
| chr5 | 30383287 | Amillepora08507 | Ras-related protein Rab-18-B | 3-prime-UTR | |
| chr5 | 30384792 | Intergenic region | --- | | |
| chr5 | 30618427 | Amillepora08533 | Double-stranded RNA-specific editase 1 | 3-prime-UTR | |
| chr5 | 30621830 | Intergenic region | Acropora digitifera RNA polymerase-associated protein LEO1-like | | 0 |
| chr5 | 30628493 | Amillepora08534 | RNA polymerase-associated protein LEO1 | 3-prime-UTR | |
| chr5 | 30644028 | Intergenic region | --- | | |
| chr5 | 30895500 | Amillepora08567 | 60S ribosomal protein L27 | Exon, AA change: Asp31Glu | |
| chr6 | 197290 | Amillepora18692 | Coadhesin | 3-prime-UTR | |
| chr6 | 835657 | Intergenic region | Acropora millepora delta-1-pyrroline-5-carboxylate dehydrogenase, mitochondrial-like | | 0 |
| chr6 | 1563742 | Amillepora18800 | Acropora millepora uncharacterized LOC114967501 | 3-prime-UTR | 0 |
| chr6 | 1730605 | Intergenic region | Acropora millepora transmembrane protein 177-like | | 2.00E-43 |
| chr6 | 2067003 | Amillepora18850 | Glutathione S-transferase kappa 1 | 3-prime-UTR | |
| chr6 | 2414402 | Intergenic region | Acropora digitifera uncharacterized | | 6.00E-38 |
| chr6 | 2414426 | Intergenic region | Acropora digitifera uncharacterized | | 6.00E-21 |
| chr6 | 3074426 | Intergenic region | Acropora digitifera uncharacterized LOC107328311 | | 0 |
| chr6 | 3246114 | Intergenic region | Acropora millepora acidic leucine-rich nuclear phosphoprotein 32 family member B-like | | 0 |
| chr6 | 3323171 | Amillepora18958 | Integrin-linked kinase-associated serine/threonine phosphatase 2C | 3-prime-UTR | |
| chr6 | 3901771 | Amillepora19013 | Endothelin-converting enzyme homolog | Exon, AA change: Glu398Gly | |
| chr6 | 4139854 | Amillepora19035 | Lens fiber membrane intrinsic protein | 3-prime-UTR | |
| chr6 | 6338568 | Amillepora19218 | Polyhomeotic-like protein 1 | 3-prime-UTR | |
| chr6 | 7820892 | Intergenic region | Acropora millepora cationic amino acid transporter 2-like | | 6.00E-60 |
| chr6 | 7882296 | Amillepora19345 | Tyrosine-protein phosphatase non-receptor type 11 | Exon synonymous variant | |
| chr6 | 8084352 | Amillepora19361 | Translocon-associated protein subunit gamma | synonymous | |
| chr6 | 8119295 | Amillepora19366 | Acropora millepora uncharacterized LOC114954782 | 3-prime-UTR | 0 |
| chr6 | 8135438 | Intergenic region | Acropora millepora myosin-11-like | | 0 |
| chr6 | 8955263 | Amillepora19435 | Acetyl-coenzyme A transporter 1 | synonymous | |
| chr6 | 9320947 | Intergenic region | --- | | |
| chr6 | 9320976 | Intergenic region | --- | | |
| chr6 | 9386595 | Intergenic region | Acropora digitifera UBX domain-containing protein 11-like | | 0 |
| chr6 | 9541508 | Amillepora19484 | Zinc finger protein 570 | Exon, AA change: Pro665Ser | |
| chr6 | 10253000 | Amillepora19558 | Profilin-4 | 3-prime-UTR | |
| chr6 | 10254355 | Intergenic region | Acropora millepora sialomucin core protein 24-like | | 0 |
| chr6 | 12291804 | Amillepora19737 | Acropora millepora uncharacterized LOC114954993 | 3-prime-UTR | 0 |
| chr6 | 12454783 | Intergenic region | Acropora millepora polycomb protein SCMH1-like | | 0 |
| chr6 | 12686972 | Intergenic region | Acropora millepora transcription factor IIIA-like | | 4.00E-25 |
| chr6 | 14109752 | Amillepora19881 | SH3 domain-containing kinase-binding protein 1 | synonymous | |
| chr6 | 14723880 | Amillepora19940 | Vacuolar protein sorting-associated protein 13C | 3-prime-UTR | |
| chr6 | 15103734 | Intergenic region | --- | | |
| chr6 | 15219742 | Intergenic region | Acropora millepora androglobin-like | | 0 |
| chr6 | 15487629 | Intergenic region | Acropora digitifera structural maintenance of chromosomes protein 6-like | | 0 |
| chr6 | 16081073 | Amillepora20049 | ADP-ribosylation factor-like protein 13B | synonymous | |
| chr6 | 16278909 | Amillepora20065 | Tyrosine--tRNA ligase, cytoplasmic | synonymous | |
| chr6 | 16840727 | Amillepora20112 | Cystatin-B | synonymous | |
| chr6 | 16910540 | Intergenic region | Acropora digitifera uncharacterized LOC107336362 | | 0 |
| chr6 | 16910903 | Intergenic region | Acropora digitifera uncharacterized LOC107336362 | | 0 |
| chr6 | 16910996 | Intergenic region | Acropora digitifera uncharacterized LOC107336362 | | 0 |
| chr6 | 17597271 | Intergenic region | Acropora digitifera baculoviral IAP repeat-containing protein 3-like | | 0 |
| chr6 | 17905638 | Intergenic region | Acropora millepora wiskott-Aldrich syndrome protein family member 3-like | | 2.00E-99 |
| chr6 | 19096683 | Intergenic region | Acropora millepora G-protein coupled receptor 161-like | | 0 |
| chr6 | 19169457 | Amillepora20298 | Alkaline phosphatase, tissue-nonspecific isozyme | Intron | |
| chr6 | 19564888 | Amillepora20323 | Probable U3 small nucleolar RNA-associated protein 11 | Exon, AA change: Arg132Cys | |
| chr6 | 19763244 | Amillepora20340 | Phosphatidylinositol-binding clathrin assembly protein | 3-prime-UTR | |
| chr7 | 13715 | Amillepora10917 | Acropora millepora uncharacterized LOC114955163 | synonymous | 0 |
| chr7 | 117090 | Amillepora10921 | Collagen alpha-1(XXVII) chain B | Exon synonymous variant | |
| chr7 | 613451 | Intergenic region | Acropora millepora IQ calmodulin-binding motif-containing protein 1-like | | 0 |
| chr7 | 720712 | Intergenic region | Acropora digitifera nucleotide-binding oligomerization domain-containing protein 2-like | | 1.00E-164 |
| chr7 | 1051388 | Amillepora10994 | ADP-ribosylation factor-like protein 5B | 3-prime-UTR | |
| chr7 | 2005721 | Intergenic region | Acropora millepora spondin-1-like | | 0 |
| chr7 | 2267278 | Amillepora11064 | THUMP domain-containing protein 1 | Exon synonymous variant | |
| chr7 | 2267327 | Amillepora11064 | THUMP domain-containing protein 1 | Exon, AA change: Val292Leu | |
| chr7 | 2267395 | Amillepora11064 | THUMP domain-containing protein 1 | 3-prime-UTR | |
| chr7 | 2267428 | Amillepora11064 | THUMP domain-containing protein 1 | 3-prime-UTR | |
| chr7 | 2274879 | Intergenic region | --- | | |
| chr7 | 2564181 | Amillepora11097 | DnaJ homolog subfamily B member 6 | 3-prime-UTR | |
| chr7 | 2675305 | Amillepora11105 | Rac GTPase-activating protein 1 | 3-prime-UTR | |
| chr7 | 4773865 | Intergenic region | Acropora millepora uncharacterized LOC114952530 | | 0 |
| chr7 | 5282232 | Intergenic region | Acropora millepora GATOR complex protein MIOS-A-like | | 1.00E-51 |
| chr7 | 5860356 | Amillepora11319 | Peptidyl-prolyl cis-trans isomerase G | synonymous | |
| chr7 | 6297850 | Amillepora11351 | EF-hand calcium-binding domain-containing protein 1; | 3-prime-UTR | |
| chr7 | 6303680 | Amillepora11352 | Protein AF-10 | 3-prime-UTR | |
| chr7 | 10647290 | Intergenic region | Acropora millepora 39S ribosomal protein L32, mitochondrial-like | | 0 |
| chr7 | 10653861 | Intergenic region | Acropora millepora transmembrane and ubiquitin-like domain-containing protein 1 | | 0 |
| chr7 | 13426518 | Intergenic region | --- | | |
| chr7 | 16088814 | Amillepora12203 | Formin-2 | 3-prime-UTR | |
| chr7 | 16515663 | Amillepora12241 | E3 ubiquitin-protein ligase MARCH9 | 3-prime-UTR | |
| chr7 | 17905193 | Amillepora12367 | Serine/threonine-protein kinase tousled-like 2 | 3-prime-UTR | |
| chr7 | 18848971 | Amillepora12460 | Integrin alpha-9 | Intron | |
| chr7 | 19616444 | Amillepora12529 | Collagen alpha-2(I) chain | 3-prime-UTR | |
| chr7 | 19616541 | Amillepora12529 | Collagen alpha-2(I) chain | 3-prime-UTR | |
| chr7 | 19966205 | Amillepora12560 | N-chimaerin | synonymous | |
| chr7 | 20526645 | Amillepora12602 | Acropora millepora uncharacterized LOC122960901 | 3-prime-UTR | 7.00E-98 |
| chr7 | 20579006 | Intergenic region | Acropora millepora vegetative incompatibility protein HET-E-1-like | | 0 |
| chr7 | 20722422 | Amillepora12614 | Acropora millepora uncharacterized LOC114955234 | 3-prime-UTR | 0 |
| chr7 | 21530249 | Amillepora12687 | Reversion-inducing cysteine-rich protein with Kazal motifs | 3-prime-UTR | |
| chr7 | 21954003 | Amillepora12716 | CCAAT/enhancer-binding protein zeta | synonymous | |
| chr7 | 22094001 | Amillepora12729 | DBH-like monooxygenase protein 1 | 3-prime-UTR | |
| chr7 | 22192821 | Amillepora12739 | Fibropellin-1 | Exon, AA change: Val660Ile | |
| chr7 | 22421606 | Amillepora12757 | DNA topoisomerase 1 | Intron | |
| chr7 | 23048433 | Amillepora12810 | CUB and peptidase domain-containing protein 1 | Exon synonymous variant | |
| chr7 | 23758462 | Amillepora12861 | Serine/threonine-protein phosphatase 6 regulatory ankyrin repeat subunit B | Intron | |
| chr8 | 961705 | Intergenic region | Acropora millepora ZP domain-containing protein-like | | 0 |
| chr8 | 2035534 | Amillepora23640 | Acropora digitifera leucine-rich repeats and immunoglobulin-like domains protein 1 | Exon, AA change: Gln82Lys | 1.00E-178 |
| chr8 | 2070152 | Amillepora23642 | Chymotrypsinogen 2 | 3-prime-UTR | |
| chr8 | 2769223 | Intergenic region | Acropora millepora uncharacterized LOC114963047 | | 0 |
| chr8 | 6487117 | Amillepora23956 | Beta-galactosidase-1-like protein 3 | 3-prime-UTR | |
| chr8 | 7487940 | Intergenic region | Acropora digitifera uncharacterized LOC107341263 | | 0 |
| chr8 | 7724188 | Amillepora24045 | FUN14 domain-containing protein 1B | 5-prime-UTR | |

| | | | | | |
|---|---|---|---|---|---|
| chr8 | 8060122 | Intergenic region | Acropora millepora uncharacterized LOC114971375 | | 0 |
| chr8 | 8538570 | Intergenic region | Acropora millepora trace amine-associated receptor 9-like | | 2.00E-161 |
| chr8 | 8668167 | Amillepora24111 | Glypican-5 | 3-prime-UTR | |
| chr8 | 8670243 | Intergenic region | --- | | |
| chr8 | 10234328 | Amillepora24234 | B-cell lymphoma 3 protein | Intron | |
| chr8 | 10289128 | Amillepora24238 | Acropora millepora ubiquitin carboxyl-terminal hydrolase 2-like | 3-prime-UTR | 0 |
| chr8 | 11483288 | Amillepora24308 | THAP domain-containing protein 4 | 3-prime-UTR | |
| chr8 | 11628941 | Amillepora24324 | Rho guanine nucleotide exchange factor 12 | 5_prime_UTR | |
| chr8 | 11887309 | Amillepora24341 | SPRY domain-containing protein 7 | 3-prime-UTR | |
| chr8 | 12066965 | Amillepora24356 | GRAM domain-containing protein 1B; | Intron | |
| chr8 | 12656822 | Amillepora24396 | Renin receptor | synonymous | |
| chr8 | 14546562 | Amillepora24553 | E3 ubiquitin-protein ligase CBL-B | Intron | |
| chr8 | 15006438 | Amillepora24584 | WASH complex subunit 2 | Exon, AA change: Pro1232Ala | |
| chr8 | 15195131 | Amillepora24597 | Major facilitator superfamily domain-containing protein 1 | 3-prime-UTR | |
| chr8 | 15861361 | Intergenic region | Acropora millepora protein BTG2-like | | 0 |
| chr8 | 15995440 | Amillepora24662 | Disks large 1 tumor suppressor protein | 3-prime-UTR | |
| chr8 | 16011821 | Intergenic region | Acropora digitifera transcription initiation factor TFIID subunit 9B-like | | 0 |
| chr8 | 16040876 | Amillepora24667 | Acropora millepora myosin-2 heavy chain-like | 5_prime_UTR | 0 |
| chr8 | 16174692 | Intergenic region | Acropora millepora isolate LDH lactate dehydrogenase-like | | 5.00E-95 |
| chr8 | 16512018 | Amillepora24719 | Integrator complex subunit 4 | synonymous | |
| chr8 | 16512397 | Intergenic region | Acropora millepora cilia- and flagella-associated protein 300-like | | 7.00E-28 |
| chr8 | 16512527 | Intergenic region | Acropora millepora cilia- and flagella-associated protein 300-like | | 7.00E-28 |
| chr8 | 16739540 | Amillepora24737 | Cilia- and flagella-associated protein 44 | synonymous | |
| chr8 | 16762086 | Intergenic region | Acropora millepora sodium/potassium-transporting ATPase subunit beta-1-like | | 0 |
| chr8 | 16920657 | Intergenic region | --- | | |
| chr8 | 17464285 | Amillepora24820 | Bromodomain testis-specific protein | Exon, AA change: Asp833Gly | |
| chr8 | 17471085 | Intergenic region | Acropora millepora dnaJ homolog subfamily C member 21-like | | 4.00E-174 |
| chr8 | 17472729 | Intergenic region | Acropora millepora dnaJ homolog subfamily C member 21-like | | 4.00E-12 |
| chr8 | 17478647 | Intergenic region | Acropora millepora VID27-like protein | | 0 |
| chr8 | 17478670 | Intergenic region | Acropora millepora VID27-like protein | | 0 |
| chr8 | 17703640 | Amillepora24846 | Stress-associated endoplasmic reticulum protein 2 | 5_prime_UTR | |
| chr9 | 704984 | Amillepora20424 | Acropora millepora uncharacterized LOC114961230 | synonymous | 0 |
| chr9 | 1252006 | Intergenic region | Acropora millepora MAM and LDL-receptor class A domain-containing protein 1-like | | 0 |
| chr9 | 1992015 | Amillepora20538 | Protein scribble homolog | 5-prime-UTR | |
| chr9 | 4644562 | Intergenic region | Acropora millepora 60S ribosomal protein L7-like | | 2.00E-60 |
| chr9 | 4644736 | Intergenic region | Acropora millepora 60S ribosomal protein L7-like | | 2.00E-60 |
| chr9 | 4996223 | Intergenic region | Acropora millepora adenosine receptor A2b-like | | 0 |
| chr9 | 6097335 | Amillepora20947 | Ras-related GTP-binding protein C | synonymous | |
| chr9 | 7364497 | Intergenic region | Acropora millepora uncharacterized LOC122962316 | | 0 |
| chr9 | 7956876 | Amillepora21104 | Bone morphogenetic protein 7 | 3-prime-UTR | |
| chr9 | 9507270 | Intergenic region | Acropora millepora deoxyhypusine hydroxylase-like | | 0 |
| chr9 | 9592013 | Intergenic region | Acropora millepora polyadenylate-binding protein 4-like | | 0 |
| chr9 | 11305321 | Intergenic region | Orbicella faveolata uncharacterized LOC110054496 | | 6.00E-50 |
| chr9 | 11539399 | Amillepora21357 | AP2-associated protein kinase 1 | 3-prime-UTR | |
| chr9 | 11907955 | Intergenic region | Acropora millepora trace amine-associated receptor 2-like | | 0 |
| chr9 | 12019152 | Amillepora21385 | Extracellular sulfatase Sulf-1 | Exon, AA change: Arg448Gln | |
| chr9 | 12503975 | Intergenic region | Acropora millepora cubilin-like | | 0 |
| chr9 | 13062554 | Intergenic region | Acropora digitifera putative nuclease HARBI1 | | 2.00E-49 |
| chr9 | 15374929 | Amillepora21577 | E3 ubiquitin-protein ligase MIB2 | Intron | |
| chr9 | 15374956 | Amillepora21577 | E3 ubiquitin-protein ligase MIB2 | Intron | |
| chr10 | 650375 | Amillepora16821 | ETS domain-containing protein Elk-1 | 5_prime_UTR | |
| chr10 | 1109095 | Amillepora16854 | Alpha-aminoadipic semialdehyde synthase, mitochondria | Exon, AA change: Thr881Ser | |
| chr10 | 2046401 | Intergenic region | Acropora millepora exportin-T-like | | 1.00E-22 |
| chr10 | 2113724 | Intergenic region | Acropora millepora uncharacterized LOC114966078 | | 5.00E-135 |
| chr10 | 2256741 | Amillepora16947 | S-adenosylmethionine sensor upstream of mTORC1 | 3-prime-UTR | |
| chr10 | 2668973 | Intergenic region | Acropora millepora uncharacterized LOC114969063 | | 2.00E-112 |
| chr10 | 3080731 | Amillepora17031 | Inactive histone-lysine N-methyltransferase 2E | Exon, AA change: Pro333Ala | |
| chr10 | 3085598 | Amillepora17031 | Inactive histone-lysine N-methyltransferase 2E | Exon, AA change: Ala84Val | |
| chr10 | 3335215 | Intergenic region | Acropora millepora uncharacterized LOC122963252 | | 0 |
| chr10 | 3840709 | Amillepora17077 | Cilia- and flagella-associated protein 100 | synonymous | |
| chr10 | 3951267 | Amillepora17089 | 25S rRNA (cytosine-C(5))-methyltransferase rcm1 | 3-prime-UTR | |
| chr10 | 10055062 | Amillepora17759 | Acropora millepora uncharacterized LOC114960967 | 3-prime-UTR | 0 |
| chr10 | 10055824 | Amillepora17759 | Acropora millepora uncharacterized LOC114960967 | 3-prime-UTR | 0 |
| chr10 | 10678205 | Amillepora17811 | Acropora millepora golgin subfamily A member 6-like protein 22 | Exon, AA change: Gln163Glu | 4.00E-111 |
| chr10 | 11478380 | Intergenic region | Acropora digitifera uncharacterized LOC107350321 | | 6.00E-112 |
| chr10 | 11754427 | Intergenic region | --- | | |
| chr10 | 11928409 | Amillepora17925 | Transcription factor HES-4 | 5-prime-UTR | |
| chr10 | 11970210 | Intergenic region | --- | | |
| chr10 | 12126851 | Amillepora17937 | Fibroblast growth factor receptor | Intron | |
| chr10 | 12396573 | Amillepora17953 | Apoptosis regulator R1 | 3-prime-UTR | |
| chr10 | 14657390 | Amillepora18085 | Fibroblast growth factor receptor 1 | 3-prime-UTR | |
| chr10 | 14696186 | Amillepora18087 | SMC5-SMC6 complex localization factor protein 2 | Exon, AA change: His86Tyr | |
| chr10 | 15950011 | Amillepora18188 | Ubiquitin-conjugating enzyme E2 L3 | 3-prime-UTR | |
| chr10 | 16412361 | Amillepora18241 | Nucleolar GTP-binding protein 2 | Exon, AA change: Asn525Ser | |
| chr10 | 16686800 | Amillepora18259 | Tubulin alpha-1C chain | synonymous | |
| chr10 | 17592201 | Intergenic region | Acropora millepora acylamino-acid-releasing enzyme-like | | 1.00E-176 |
| chr10 | 17964704 | Amillepora18353 | Putative helicase mov-10-B.1 | Exon, AA change: Ser102Asn | |
| chr10 | 19125756 | Amillepora18462 | ATP-dependent RNA helicase MAK5 | synonymous | |
| chr10 | 19220843 | Intergenic region | Acropora millepora neuromedin-K receptor-like | | 0 |
| chr10 | 19790714 | Amillepora18526 | Sodium/glucose cotransporter 5 | Exon, AA change: Ile647Leu | |
| chr10 | 19800682 | Intergenic region | Acropora millepora sodium/glucose cotransporter 5-like | | 1.00E-164 |
| chr10 | 20035791 | Amillepora18543 | Coatomer subunit gamma-2 | synonymous | |
| chr11 | 736523 | Amillepora26329 | 60S ribosomal protein L10 | 5-prime-UTR | |
| chr11 | 1825060 | Amillepora26477 | Protein SMG5 | 3-prime-UTR | |
| chr11 | 2375120 | Amillepora26519 | DnaJ homolog subfamily C member 10 | 5_prime_UTR | |
| chr11 | 2740588 | Amillepora26549 | Protein phosphatase 1 regulatory subunit 21 | Exon, AA change: Val99Met | |
| chr11 | 3174940 | Amillepora26581 | Homeobox protein Nkx-2.2a | 3-prime-UTR | |
| chr11 | 3389866 | Intergenic region | Acropora millepora pre-mRNA-splicing factor CWC25 homolog | | 0 |
| chr11 | 3686390 | Amillepora26623 | 1-phosphatidylinositol 4,5-bisphosphate phosphodiesterase beta-4; | synonymous | |
| chr11 | 4194006 | Intergenic region | Acropora millepora endosome/lysosome-associated apoptosis and autophagy regulator family member 2-like | | 0 |
| chr11 | 4744260 | Intergenic region | Acropora millepora adenylosuccinate lyase-like | | 5.00E-51 |
| chr11 | 5856944 | Amillepora26778 | Multifunctional methyltransferase subunit TRM112-like protein | 3-prime-UTR | |
| chr11 | 11062659 | Intergenic region | Actinia tenebrosa dynein assembly factor 3, axonemal-like | | 2.00E-36 |
| chr11 | 11654242 | Amillepora27228 | Kalirin | Intron | |
| chr11 | 12260874 | Amillepora27264 | Heterogeneous nuclear ribonucleoprotein U-like protein 1 | Intron | |
| chr11 | 13830803 | Amillepora27358 | 26S proteasome non-ATPase regulatory subunit 7 | 3-prime-UTR | |
| chr11 | 13959481 | Amillepora27376 | Dual adapter for phosphotyrosine and 3-phosphotyrosine and 3-phosphoinositide | 3-prime-UTR | |
| chr11 | 14111681 | Amillepora27386 | Beta-1,3-galactosyltransferase 6 | Exon, AA change: Met322Arg | |
| chr11 | 14842966 | Amillepora27445 | Homeobox protein OTX1 B | synonymous | |
| chr11 | 15527695 | Amillepora27497 | Serine/threonine-protein kinase AtPK1/AtPK6 | Exon, AA change: Ala1026Thr | |
| chr11 | 15748771 | Intergenic region | Acropora digitifera uncharacterized LOC107337576 | | 0 |
| chr12 | 1372976 | Amillepora13039 | Toll-like receptor 13 | 3-prime-UTR | |
| chr12 | 1885226 | Amillepora13086 | --- | Exon, AA change: Val8Phe | |
| chr12 | 2196516 | Amillepora13107 | Tetraspanin-11 | 3-prime-UTR | |
| chr12 | 2899750 | Amillepora13161 | Zinc finger protein 888 | 5_prime_UTR | |
| chr12 | 2951021 | Amillepora13169 | Acropora digitifera RAD51-associated protein 1-like | 3-prime-UTR | 0 |
| chr12 | 3362733 | Intergenic region | Acropora digitifera protein EFR3 homolog B-like | | 2.00E-49 |
| chr12 | 3665877 | Intergenic region | Acropora digitifera spermatogenesis-defective protein 39 homolog | | 2.00E-31 |
| chr12 | 3699900 | Amillepora13239 | -- | 3-prime-UTR | |
| chr12 | 3762566 | Amillepora13244 | Low-density lipoprotein receptor-related protein 6 | synonymous | |
| chr12 | 4793703 | Amillepora13327 | Acropora millepora serine/arginine repetitive matrix protein 2-like | 3-prime-UTR | 0 |
| chr12 | 4795986 | Amillepora13327 | Acropora millepora serine/arginine repetitive matrix protein 2-like | 3-prime-UTR | 0 |
| chr12 | 6711950 | Amillepora13525 | RNA polymerase-associated protein CTR9 homolog | synonymous | |
| chr12 | 6803700 | Amillepora13533 | ZP domain-containing protein | 3-prime-UTR | |
| chr12 | 8179454 | Intergenic region | Acropora millepora F-box/LRR-repeat protein 20-like | | 0 |
| chr12 | 8206891 | Intergenic region | Acropora millepora uncharacterized LOC114948244 | | 3.00E-79 |
| chr12 | 9110857 | Amillepora13721 | Sodium-independent sulfate anion transporter | intron | |
| chr12 | 10368054 | Amillepora13838 | Acropora digitifera uncharacterized LOC107355650 | intron | 0 |
| chr12 | 12357206 | Amillepora13992 | Cysteine-rich motor neuron 1 protein | 3-prime-UTR | |
| chr12 | 13161748 | Amillepora14061 | Acropora millepora uncharacterized LOC114971494 | 3-prime-UTR | 0 |
| chr12 | 14265972 | Amillepora14127 | Anoctamin-4 | 3-prime-UTR | |
| chr12 | 15872734 | Amillepora14260 | Rho GTPase-activating protein 44 | 3-prime-UTR | |
| chr12 | 17827684 | Amillepora14418 | Probable E3 ubiquitin-protein ligase MGRN1 | 3-prime-UTR | |
| chr12 | 17827711 | Amillepora14418 | Probable E3 ubiquitin-protein ligase MGRN1 | 3-prime-UTR | |
| chr12 | 18282402 | Intergenic region | Acropora millepora ketimine reductase mu-crystallin-like | | 5.00E-160 |
| chr12 | 18282411 | Intergenic region | Acropora millepora ketimine reductase mu-crystallin-like | | 5.00E-160 |
| chr12 | 18282920 | Amillepora14462 | Ketimine reductase mu-crystallin | 3-prime-UTR | |
| chr12 | 18609890 | Intergenic region | Acropora millepora myosin heavy chain, striated muscle-like | | 0 |
| chr12 | 18864390 | Intergenic region | Acropora millepora polycystic kidney disease protein 1-like 2 | | 0 |
| chr12 | 19357487 | Amillepora14546 | Hemagglutinin/amebocyte aggregation factor | Exon, AA change: Pro260Arg | |
| chr12 | 20403989 | Amillepora14626 | Dolichyl-diphosphooligosaccharide--protein glycosyltransferase subunit STT3A | synonymous | |

| | | | | | |
|---|---|---|---|---|---|
| chr12 | 20969631 | Amillepora14686 | Acropora millepora uncharacterized LOC114957676 | Intron | 0 |
| chr12 | 21125848 | Intergenic region | Acropora millepora coiled-coil domain-containing protein 137-like | | 0 |
| chr12 | 21800943 | Intergenic region | Acropora digitifera uncharacterized LOC107335474 | | 0 |
| chr12 | 21863110 | Amillepora14742 | Regulatory-associated protein of mTOR | Intron | |
| chr12 | 22196411 | Amillepora14764 | Aryl hydrocarbon receptor | 3-prime-UTR | |
| chr12 | 22196562 | Amillepora14764 | Aryl hydrocarbon receptor | 3-prime-UTR | |
| chr12 | 22259695 | Intergenic region | Acropora millepora tropomyosin-like | | 0 |
| chr12 | 22403792 | Amillepora14777 | ATP-binding cassette sub-family A member 5 | 3-prime-UTR | |
| chr12 | 22405818 | Intergenic region | Acropora millepora uncharacterized | | 1.00E-61 |
| chr12 | 22458014 | Intergenic region | Acropora millepora dynactin subunit 5-like | | 0 |
| chr12 | 23022357 | Amillepora14835 | ATP-binding cassette sub-family A member 3 | 3-prime-UTR | |
| chr13 | 41863 | Amillepora14866 | Protein-glucosylgalactosylhydroxylysine glucosidase | Exon, AA change: Ser1184Phe | |
| chr13 | 90546 | Amillepora14876 | N-terminal kinase-like protein | synonymous | |
| chr13 | 157206 | Intergenic region | Acropora millepora synaptotagmin-5-like | | 0 |
| chr13 | 1844351 | Intergenic region | Acropora millepora uncharacterized LOC114956613 | | 0 |
| chr13 | 2758517 | Amillepora15111 | Serine/threonine-protein phosphatase 2A regulatory subunit B'' subunit gamma | 3-prime-UTR | |
| chr13 | 3224179 | Amillepora15159 | DNA-directed RNA polymerase III subunit RPC6 | synonymous | |
| chr13 | 3291660 | Intergenic region | Acropora millepora putative ATP-dependent RNA helicase DHX57 | | 0 |
| chr13 | 3768600 | Amillepora15207 | Mammalian ependymin-related protein 1 | Exon, AA change: Val5Leu | |
| chr13 | 4891673 | Amillepora15292 | Poly [ADP-ribose] polymerase 1 | synonymous | |
| chr13 | 5205616 | Amillepora15323 | Cardiolipin synthase (CMP-forming) | synonymous | |
| chr13 | 5211883 | Intergenic region | Acropora millepora uncharacterized | | 4.00E-88 |
| chr13 | 5557600 | Intergenic region | --- | | |
| chr13 | 7330152 | Intergenic region | Acropora millepora uncharacterized LOC114954830 | | 0 |
| chr13 | 7549237 | Amillepora15521 | COUP transcription factor 2 | 3-prime-UTR | |
| chr13 | 7916532 | Amillepora15554 | Cholesterol 24-hydroxylase | synonymous | |
| chr13 | 8062598 | Intergenic region | Acropora millepora protein NDRG3-like | | 0 |
| chr13 | 10517892 | Amillepora15763 | Transmembrane protein Tmp21 | 3-prime-UTR | |
| chr13 | 10518356 | Amillepora15763 | Transmembrane protein Tmp21 | 3-prime-UTR | |
| chr13 | 10706805 | Intergenic region | Acropora millepora ubiquitin carboxyl-terminal hydrolase 34-like | | 0 |
| chr13 | 12726423 | Amillepora16017 | Mediator of RNA polymerase II transcription subunit 14 | 3-prime-UTR | |
| chr13 | 13476017 | Amillepora16062 | Inositol-trisphosphate 3-kinase B | 3-prime-UTR | |
| chr13 | 13573035 | Intergenic region | Acropora millepora uncharacterized | | 4.00E-115 |
| chr13 | 13633555 | Amillepora16078 | Annexin A13 | Exon, AA change: Ala212Val | |
| chr13 | 13935406 | Amillepora16106 | COMM domain-containing protein 6 | 3-prime-UTR | |
| chr13 | 14813047 | Intergenic region | Acropora millepora uncharacterized | | 7.00E-58 |
| chr13 | 15001451 | Intergenic region | Acropora millepora uncharacterized LOC114960791 | | 0 |
| chr13 | 15229631 | Intergenic region | Acropora millepora uncharacterized LOC114960773 | | 0 |
| chr13 | 15853694 | Intergenic region | Acropora millepora polyribonucleotide nucleotidyltransferase 1, mitochondrial-like | | 7.00E-175 |
| chr13 | 15853842 | Intergenic region | Acropora millepora polyribonucleotide nucleotidyltransferase 1, mitochondrial-like | | 7.00E-175 |
| chr13 | 15853888 | Intergenic region | Acropora millepora polyribonucleotide nucleotidyltransferase 1, mitochondrial-like | | 7.00E-175 |
| chr13 | 16025681 | Amillepora16259 | Acropora millepora uncharacterized LOC114946805 | 3-prime-UTR | 0 |
| chr13 | 16344052 | Amillepora16298 | RNA polymerase-associated protein RTF1 homolog | synonymous | |
| chr13 | 17126354 | Amillepora16358 | Zinc finger RNA-binding protein | 3-prime-UTR | |
| chr13 | 18312643 | Intergenic region | Acropora millepora zinc phosphodiesterase ELAC protein 1-like | | 0 |
| chr13 | 18540764 | Intergenic region | --- | | |
| chr13 | 18560737 | Amillepora16446 | Glycine N-methyltransferase | 3-prime-UTR | |
| chr13 | 19049062 | Amillepora16476 | Equistatin | Exon, AA change: Gly533Ala | |
| chr13 | 19105504 | Amillepora16483 | Acropora millepora uncharacterized LOC114956812 | Exon, AA change: Ile273Val | 0 |
| chr13 | 19300613 | Intergenic region | --- | | |
| chr13 | 19779798 | Amillepora16553 | T-complex protein 1 subunit delta | Exon, AA change: Ala512Val | |
| chr13 | 20537229 | Amillepora16628 | Mediator of RNA polymerase II transcription subunit 29 | 3-prime-UTR | |
| chr13 | 20537305 | Amillepora16628 | Mediator of RNA polymerase II transcription subunit 29 | 3-prime-UTR | |
| chr13 | 20967041 | Intergenic region | Acropora digitifera inactive tyrosine-protein kinase 7-like | | 0 |
| chr13 | 21052181 | Amillepora16669 | Zinc finger protein DPF3 | 3-prime-UTR | |
| chr13 | 21663628 | Intergenic region | Acropora millepora uncharacterized LOC114957813 | | 0 |
| chr13 | 21748705 | Amillepora16717 | Kinesin-like protein KIF16B | 3-prime-UTR | |
| chr13 | 21749500 | Intergenic region | Acropora millepora serine protease 23-like | | 0 |
| chr14 | 588168 | Amillepora24928 | Serine/arginine-rich splicing factor 1 | 3-prime-UTR | |
| chr14 | 608757 | Amillepora24931 | Serine/threonine-protein phosphatase PP1-beta | 3-prime-UTR | |
| chr14 | 946793 | Amillepora24967 | Small G protein signaling modulator 2 | 3-prime-UTR | |
| chr14 | 948315 | Amillepora24967 | Small G protein signaling modulator 2 | Splicing variant | |
| chr14 | 953049 | Amillepora24967 | Small G protein signaling modulator 2 | Exon, AA change: Thr287Ile | |
| chr14 | 1055527 | Amillepora24982 | SH3 and PX domain-containing protein 2B | 3-prime-UTR | |
| chr14 | 1108670 | Intergenic region | Acropora millepora apoptosis-inducing factor 3-like | | 0 |
| chr14 | 1108818 | Amillepora24987 | Apoptosis-inducing factor 3 | 3-prime-UTR | |
| chr14 | 1118862 | Amillepora24988 | Cytochrome c oxidase subunit 6A, mitochondrial | synonymous | |
| chr14 | 1315559 | Amillepora25017 | Protein phosphatase 1D | 3-prime-UTR | |
| chr14 | 1596904 | Intergenic region | Acropora millepora tumor protein 63-like | | 1.00E-76 |
| chr14 | 1907353 | Amillepora25070 | Histone-lysine N-methyltransferase, H3 lysine-79 specific | 3-prime-UTR | |
| chr14 | 1907486 | Amillepora25070 | Histone-lysine N-methyltransferase, H3 lysine-79 specific | 3-prime-UTR | |
| chr14 | 2002620 | Amillepora25075 | Importin-13 | 3-prime-UTR | |
| chr14 | 2152350 | Intergenic region | Acropora millepora transcription factor AP-1-like | | 0 |
| chr14 | 3581047 | Intergenic region | Acropora millepora uncharacterized LOC114968405 | | 3.00E-103 |
| chr14 | 3688161 | Amillepora25225 | Inactive tyrosine-protein kinase transmembrane receptor ROR1 | Intron | |
| chr14 | 3896603 | Amillepora25238 | Tubulin polyglutamylase TTLL7 | 3-prime-UTR | |
| chr14 | 4051136 | Intergenic region | Acropora millepora trithorax group protein osa-like | | 0 |
| chr14 | 4777188 | Amillepora25320 | E3 ubiquitin-protein ligase DZIP3 | Exon, AA change: Ser182Phe | |
| chr14 | 4837771 | Amillepora25325 | Epidermal growth factor receptor substrate 15-like 1 | Exon, AA change: His78Arg | |
| chr14 | 4962100 | Amillepora25339 | Cytochrome c-type heme lyase | 3-prime-UTR | |
| chr14 | 5371942 | Amillepora25361 | Acropora millepora uncharacterized LOC114970783 | Exon, AA change: Met59Ile | 0 |
| chr14 | 8244042 | Amillepora25596 | Rho GTPase-activating protein 29 | Intron | |
| chr14 | 12311118 | Intergenic region | Acropora digitifera uncharacterized, ncRNA | | 7.00E-112 |
| chr14 | 12829505 | Amillepora25933 | Protein YIPF1 | 3-prime-UTR | |
| chr14 | 13051006 | Intergenic region | Acropora digitifera urokinase plasminogen activator surface receptor-like | | 4.00E-66 |
| chr14 | 14163032 | Intergenic region | Dipsastraea rotumana isolate WF101 Pax-C gene | | 2.00E-70 |
| chr14 | 14350557 | Amillepora26017 | Peptidylprolyl isomerase domain and WD repeat-containing protein 1 | 3-prime-UTR | |
| chr14 | 14408091 | Amillepora26021 | Midasin | Exon, AA change: Glu617Gly | |
| chr14 | 14817113 | Intergenic region | Acropora millepora probable serine/threonine-protein kinase drkD | | 3.00E-178 |
| chr14 | 15207002 | Amillepora26077 | Receptor-type tyrosine-protein phosphatase delta | synonymous | |
| chr14 | 15207046 | Amillepora26077 | Receptor-type tyrosine-protein phosphatase delta | Exon, AA change: Ser152Gly | |
| chr14 | 15777043 | Amillepora26119 | Serine/threonine-protein kinase NIM1 | synonymous | |
| chr14 | 15867608 | Intergenic region | --- | | |
| chr14 | 16149066 | Amillepora26149 | Delta(3,5)-Delta(2,4)-dienoyl-CoA isomerase, mitochondrial | Intron | |
| chr14 | 16194833 | Amillepora26155 | Probable rRNA-processing protein EBP2 | Exon, AA change: Arg38Leu | |
| chr14 | 16208922 | Amillepora26155 | Probable rRNA-processing protein EBP2 | Exon, AA change: Thr455Pro | |
| chr14 | 16292372 | Amillepora26166 | Acropora millepora trichohyalin-like | synonymous | 0 |
| chr14 | 16300558 | Amillepora26166 | Acropora millepora trichohyalin-like | Exon, AA change: Arg191Gln | 0 |
| chr14 | 16807263 | Amillepora26202 | Acropora millepora uncharacterized LOC114963538 | Exon, AA change: Ala76Val | 7.00E-111 |
| chr14 | 17064871 | Amillepora26221 | E3 ubiquitin-protein ligase Topors | 3-prime-UTR | |
| chr14 | 17083324 | Amillepora26224 | UDP-N-acetylglucosamine transferase subunit ALG14 homolog | 3-prime-UTR | |
| chr14 | 17186023 | Amillepora26234 | Tyrosine-protein kinase SYK | Exon synonymous variant | |
| chr14 | 17186041 | Amillepora26234 | Tyrosine-protein kinase SYK | Exon synonymous variant | |
| Sc0000189 | 309672 | Amillepora33954 | Serine/threonine-protein kinase WNK1 | Intron | |
