## Supplementary Material 3 for "Synergistic genomic mechanisms of adaptation to ocean acidification in a coral holobiont"

**Supplementary Material 3. Gene Set Enrichment Analysis (GSEA) results**

| GO.ID | Term | Annotated | Significant | Pvalue | Class | gene | FDR |
|---|---|---|---|---|---|---|---|
| GO:0045603 | positive regulation of endothelial cell differentiation | 2 | 2 | 0.00039 | BP | Amillepora05687-RA, Amil | 0.220993171 |
| GO:0045621 | positive regulation of lymphocyte differentiation | 10 | 3 | 0.00083 | BP | Amillepora10499-RA, Amil | 0.220993171 |
| GO:0001711 | endodermal cell fate commitment | 3 | 2 | 0.00116 | BP | Amillepora16298-RA, Amil | 0.220993171 |
| GO:0000278 | mitotic cell cycle | 165 | 10 | 0.00159 | BP | Amillepora05723-RA, Amil | 0.220993171 |
| GO:0050685 | positive regulation of mRNA processing | 4 | 2 | 0.00229 | BP | Amillepora08533-RA, Amil | 0.220993171 |
| GO:0031440 | regulation of mRNA 3'-end processing | 4 | 2 | 0.00229 | BP | Amillepora24928-RA, Amil | 0.220993171 |
| GO:1902600 | proton transmembrane transport | 48 | 5 | 0.00249 | BP | Amillepora24987-RA, Amil | 0.220993171 |
| GO:0048468 | cell development | 237 | 12 | 0.00259 | BP | Amillepora19345-RA, Amil | 0.220993171 |
| GO:0051707 | response to other organism | 148 | 9 | 0.00266 | BP | Amillepora15159-RA, Amil | 0.220993171 |
| GO:0048871 | multicellular organismal homeostasis | 49 | 5 | 0.00273 | BP | Amillepora21104-RA, Amil | 0.220993171 |
| GO:0030336 | negative regulation of cell migration | 16 | 3 | 0.00356 | BP | Amillepora09624-RA, Amil | 0.220993171 |
| GO:0048488 | synaptic vesicle endocytosis | 5 | 2 | 0.00377 | BP | Amillepora20340-RA, Amil | 0.220993171 |
| GO:0060415 | muscle tissue morphogenesis | 5 | 2 | 0.00377 | BP | Amillepora16669-RA, Amil | 0.220993171 |
| GO:0051569 | regulation of histone H3-K4 methylation | 5 | 2 | 0.00377 | BP | Amillepora13525-RA, Amil | 0.220993171 |
| GO:0051321 | meiotic cell cycle | 34 | 4 | 0.00437 | BP | Amillepora24931-RA, Amil | 0.220993171 |
| GO:0060395 | SMAD protein signal transduction | 6 | 2 | 0.00558 | BP | Amillepora21104-RA, Amil | 0.220993171 |
| GO:0046890 | regulation of lipid biosynthetic process | 19 | 3 | 0.0059 | BP | Amillepora14835-RA, Amil | 0.220993171 |
| GO:0007010 | cytoskeleton organization | 265 | 12 | 0.00634 | BP | Amillepora05723-RA, Amil | 0.220993171 |
| GO:0046165 | alcohol biosynthetic process | 38 | 4 | 0.00654 | BP | Amillepora21104-RA, Amil | 0.220993171 |
| GO:0045785 | positive regulation of cell adhesion | 20 | 3 | 0.00684 | BP | Amillepora10499-RA, Amil | 0.220993171 |
| GO:0030595 | leukocyte chemotaxis | 7 | 2 | 0.00771 | BP | Amillepora09624-RA, Amil | 0.220993171 |
| GO:0043154 | negative regulation of cysteine-type endopeptidase activity involved in apoptotic process | 7 | 2 | 0.00771 | BP | Amillepora09624-RA, Amil | 0.220993171 |
| GO:0034067 | protein localization to Golgi apparatus | 7 | 2 | 0.00771 | BP | Amillepora15763-RA, Amil | 0.220993171 |
| GO:0072583 | clathrin-dependent endocytosis | 7 | 2 | 0.00771 | BP | Amillepora20340-RA, Amil | 0.220993171 |
| GO:0022402 | cell cycle process | 206 | 10 | 0.00779 | BP | Amillepora00093-RA, Amil | 0.220993171 |
| GO:0045582 | positive regulation of T cell differentiation | 8 | 2 | 0.01015 | BP | Amillepora10499-RA, Amil | 0.220993171 |
| GO:0006952 | defense response | 152 | 8 | 0.01062 | BP | Amillepora09624-RA, Amil | 0.220993171 |
| GO:0010243 | response to organonitrogen compound | 123 | 7 | 0.01107 | BP | Amillepora07010-RA, Amil | 0.220993171 |
| GO:0034504 | protein localization to nucleus | 45 | 4 | 0.01186 | BP | Amillepora23397-RA, Amil | 0.220993171 |
| GO:0034645 | cellular macromolecule biosynthetic process | 1168 | 34 | 0.01196 | BP | Amillepora09679-RA, Amil | 0.220993171 |
| GO:0001819 | positive regulation of cytokine production | 25 | 3 | 0.01283 | BP | Amillepora15159-RA, Amil | 0.220993171 |
| GO:0006084 | acetyl-CoA metabolic process | 9 | 2 | 0.01288 | BP | Amillepora16854-RA, Amil | 0.220993171 |
| GO:0007064 | mitotic sister chromatid cohesion | 9 | 2 | 0.01288 | BP | Amillepora08867-RA, Amil | 0.220993171 |
| GO:0050714 | positive regulation of protein secretion | 9 | 2 | 0.01288 | BP | Amillepora15763-RA, Amil | 0.220993171 |
| GO:0010033 | response to organic substance | 329 | 13 | 0.01359 | BP | Amillepora10499-RA, Amil | 0.220993171 |
| GO:0010628 | positive regulation of gene expression | 72 | 5 | 0.01394 | BP | Amillepora03480-RA, Amil | 0.220993171 |
| GO:0001816 | cytokine production | 48 | 4 | 0.0148 | BP | Amillepora15763-RA, Amil | 0.220993171 |
| GO:0001817 | regulation of cytokine production | 48 | 4 | 0.0148 | BP | Amillepora15763-RA, Amil | 0.220993171 |
| GO:0010468 | regulation of gene expression | 848 | 26 | 0.01556 | BP | Amillepora17925-RA, Amil | 0.220993171 |
| GO:0008284 | positive regulation of cell population proliferation | 49 | 4 | 0.01587 | BP | Amillepora21104-RA, Amil | 0.220993171 |
| GO:0045494 | photoreceptor cell maintenance | 10 | 2 | 0.01589 | BP | Amillepora00666-RA, Amil | 0.220993171 |
| GO:0048589 | developmental growth | 75 | 5 | 0.0164 | BP | Amillepora22668-RA, Amil | 0.220993171 |
| GO:1901698 | response to nitrogen compound | 134 | 7 | 0.0171 | BP | Amillepora07010-RA, Amil | 0.220993171 |
| GO:0098609 | cell-cell adhesion | 51 | 4 | 0.01816 | BP | Amillepora21104-RA, Amil | 0.220993171 |
| GO:0048880 | sensory system development | 77 | 5 | 0.0182 | BP | Amillepora15159-RA, Amil | 0.220993171 |
| GO:0000280 | nuclear division | 78 | 5 | 0.01914 | BP | Amillepora22273-RA, Amil | 0.220993171 |
| GO:0045580 | regulation of T cell differentiation | 11 | 2 | 0.01918 | BP | Amillepora10499-RA, Amil | 0.220993171 |
| GO:0050870 | positive regulation of T cell activation | 11 | 2 | 0.01918 | BP | Amillepora10499-RA, Amil | 0.220993171 |
| GO:0019319 | hexose biosynthetic process | 11 | 2 | 0.01918 | BP | Amillepora10499-RA, Amil | 0.220993171 |
| GO:0006094 | gluconeogenesis | 11 | 2 | 0.01918 | BP | Amillepora10499-RA, Amil | 0.220993171 |
| GO:0030865 | cortical cytoskeleton organization | 11 | 2 | 0.01918 | BP | Amillepora00093-RA, Amil | 0.220993171 |
| GO:1903039 | positive regulation of leukocyte cell-cell adhesion | 11 | 2 | 0.01918 | BP | Amillepora10499-RA, Amil | 0.220993171 |
| GO:0032868 | response to insulin | 29 | 3 | 0.01925 | BP | Amillepora02951-RA, Amil | 0.220993171 |
| GO:0022407 | regulation of cell-cell adhesion | 29 | 3 | 0.01925 | BP | Amillepora04708-RA, Amil | 0.220993171 |
| GO:0045892 | negative regulation of DNA-templated transcription | 107 | 6 | 0.01946 | BP | Amillepora21104-RA, Amil | 0.220993171 |
| GO:1905412 | negative regulation of mitotic cohesin loading | 1 | 1 | 0.01987 | BP | Amillepora21960-RA | 0.220993171 |
| GO:0021610 | facial nerve morphogenesis | 1 | 1 | 0.01987 | BP | Amillepora08533-RA | 0.220993171 |
| GO:0021618 | hypoglossal nerve morphogenesis | 1 | 1 | 0.01987 | BP | Amillepora08533-RA | 0.220993171 |
| GO:0071281 | cellular response to iron ion | 1 | 1 | 0.01987 | BP | Amillepora21104-RA | 0.220993171 |
| GO:0120186 | negative regulation of protein localization to chromatin | 1 | 1 | 0.01987 | BP | Amillepora21960-RA | 0.220993171 |
| GO:1905405 | regulation of mitotic cohesin loading | 1 | 1 | 0.01987 | BP | Amillepora21960-RA | 0.220993171 |
| GO:0071921 | cohesin loading | 1 | 1 | 0.01987 | BP | Amillepora21960-RA | 0.220993171 |
| GO:0071922 | regulation of cohesin loading | 1 | 1 | 0.01987 | BP | Amillepora21960-RA | 0.220993171 |
| GO:0071923 | negative regulation of cohesin loading | 1 | 1 | 0.01987 | BP | Amillepora21960-RA | 0.220993171 |
| GO:0050884 | neuromuscular process controlling posture | 1 | 1 | 0.01987 | BP | Amillepora08533-RA | 0.220993171 |
| GO:0090558 | plant epidermis development | 1 | 1 | 0.01987 | BP | Amillepora22668-RA | 0.220993171 |
| GO:0015876 | acetyl-CoA transport | 1 | 1 | 0.01987 | BP | Amillepora19435-RA | 0.220993171 |
| GO:0048364 | root development | 1 | 1 | 0.01987 | BP | Amillepora22668-RA | 0.220993171 |
| GO:1901337 | thioester transport | 1 | 1 | 0.01987 | BP | Amillepora19435-RA | 0.220993171 |
| GO:1903292 | protein localization to Golgi membrane | 1 | 1 | 0.01987 | BP | Amillepora10994-RA | 0.220993171 |
| GO:0043380 | regulation of memory T cell differentiation | 1 | 1 | 0.01987 | BP | Amillepora10499-RA | 0.220993171 |
| GO:0043382 | positive regulation of memory T cell differentiation | 1 | 1 | 0.01987 | BP | Amillepora10499-RA | 0.220993171 |
| GO:0048387 | negative regulation of retinoic acid receptor signaling pathway | 1 | 1 | 0.01987 | BP | Amillepora00093-RA | 0.220993171 |
| GO:0043379 | memory T cell differentiation | 1 | 1 | 0.01987 | BP | Amillepora10499-RA | 0.220993171 |
| GO:0071712 | ER-associated misfolded protein catabolic process | 1 | 1 | 0.01987 | BP | Amillepora22025-RA | 0.220993171 |
| GO:0032732 | positive regulation of interleukin-1 production | 1 | 1 | 0.01987 | BP | Amillepora15763-RA | 0.220993171 |
| GO:0019474 | L-lysine catabolic process to acetyl-CoA | 1 | 1 | 0.01987 | BP | Amillepora16854-RA | 0.220993171 |
| GO:0019477 | L-lysine catabolic process | 1 | 1 | 0.01987 | BP | Amillepora16854-RA | 0.220993171 |
| GO:0062022 | mitotic cohesin ssDNA (lagging strand) loading | 1 | 1 | 0.01987 | BP | Amillepora21960-RA | 0.220993171 |
| GO:0060717 | chorion development | 1 | 1 | 0.01987 | BP | Amillepora11097-RA | 0.220993171 |
| GO:0090627 | plant epidermal cell differentiation | 1 | 1 | 0.01987 | BP | Amillepora22668-RA | 0.220993171 |
| GO:1905245 | regulation of aspartic-type peptidase activity | 1 | 1 | 0.01987 | BP | Amillepora15763-RA | 0.220993171 |
| GO:1905246 | negative regulation of aspartic-type peptidase activity | 1 | 1 | 0.01987 | BP | Amillepora15763-RA | 0.220993171 |
| GO:0030878 | thyroid gland development | 1 | 1 | 0.01987 | BP | Amillepora05687-RA | 0.220993171 |
| GO:0022622 | root system development | 1 | 1 | 0.01987 | BP | Amillepora22668-RA | 0.220993171 |
| GO:0048486 | parasympathetic nervous system development | 1 | 1 | 0.01987 | BP | Amillepora08533-RA | 0.220993171 |
| GO:0002292 | T cell differentiation involved in immune response | 1 | 1 | 0.01987 | BP | Amillepora10499-RA | 0.220993171 |
| GO:0035964 | COPI-coated vesicle budding | 1 | 1 | 0.01987 | BP | Amillepora15763-RA | 0.220993171 |
| GO:0021965 | spinal cord ventral commissure morphogenesis | 1 | 1 | 0.01987 | BP | Amillepora08533-RA | 0.220993171 |
| GO:0002286 | T cell activation involved in immune response | 1 | 1 | 0.01987 | BP | Amillepora10499-RA | 0.220993171 |
| GO:0106111 | regulation of mitotic cohesin ssDNA (lagging strand) loading | 1 | 1 | 0.01987 | BP | Amillepora21960-RA | 0.220993171 |
| GO:0031442 | positive regulation of mRNA 3'-end processing | 1 | 1 | 0.01987 | BP | Amillepora08534-RA | 0.220993171 |
| GO:0060586 | multicellular organismal iron ion homeostasis | 1 | 1 | 0.01987 | BP | Amillepora21104-RA | 0.220993171 |
| GO:0043129 | surfactant homeostasis | 1 | 1 | 0.01987 | BP | Amillepora14835-RA | 0.220993171 |
| GO:1903867 | extraembryonic membrane development | 1 | 1 | 0.01987 | BP | Amillepora11097-RA | 0.220993171 |
| GO:0043114 | regulation of vascular permeability | 1 | 1 | 0.01987 | BP | Amillepora21104-RA | 0.220993171 |

| | | | | | | | |
|---|---|---|---|---|---|---|---|
| GO:0043117 | positive regulation of vascular permeability | 1 | 1 | 0.01987 | BP | Amillepora21104-RA | 0.220993171 |
| GO:0080147 | root hair cell development | 1 | 1 | 0.01987 | BP | Amillepora22668-RA | 0.220993171 |
| GO:0006553 | lysine metabolic process | 1 | 1 | 0.01987 | BP | Amillepora16854-RA | 0.220993171 |
| GO:0006554 | lysine catabolic process | 1 | 1 | 0.01987 | BP | Amillepora16854-RA | 0.220993171 |
| GO:0044387 | negative regulation of protein kinase activity by regulation of protein phosphorylation | 1 | 1 | 0.01987 | BP | Amillepora08533-RA | 0.220993171 |
| GO:0061402 | positive regulation of transcription from RNA polymerase II promoter in response to acidic pH | 1 | 1 | 0.01987 | BP | Amillepora10499-RA | 0.220993171 |
| GO:0071398 | cellular response to fatty acid | 1 | 1 | 0.01987 | BP | Amillepora04680-RA | 0.220993171 |
| GO:0021783 | preganglionic parasympathetic fiber development | 1 | 1 | 0.01987 | BP | Amillepora08533-RA | 0.220993171 |
| GO:0010053 | root epidermal cell differentiation | 1 | 1 | 0.01987 | BP | Amillepora22668-RA | 0.220993171 |
| GO:0010054 | trichoblast differentiation | 1 | 1 | 0.01987 | BP | Amillepora22668-RA | 0.220993171 |
| GO:0106272 | protein localization to ERGIC | 1 | 1 | 0.01987 | BP | Amillepora15763-RA | 0.220993171 |
| GO:0106273 | cytosol to ERGIC protein transport | 1 | 1 | 0.01987 | BP | Amillepora15763-RA | 0.220993171 |
| GO:2000510 | positive regulation of dendritic cell chemotaxis | 1 | 1 | 0.01987 | BP | Amillepora00093-RA | 0.220993171 |
| GO:0033700 | phospholipid efflux | 1 | 1 | 0.01987 | BP | Amillepora14835-RA | 0.220993171 |
| GO:0009085 | lysine biosynthetic process | 1 | 1 | 0.01987 | BP | Amillepora16854-RA | 0.220993171 |
| GO:0045109 | intermediate filament organization | 1 | 1 | 0.01987 | BP | Amillepora11097-RA | 0.220993171 |
| GO:0060391 | positive regulation of SMAD protein signal transduction | 1 | 1 | 0.01987 | BP | Amillepora21104-RA | 0.220993171 |
| GO:0010015 | root morphogenesis | 1 | 1 | 0.01987 | BP | Amillepora22668-RA | 0.220993171 |
| GO:2000508 | regulation of dendritic cell chemotaxis | 1 | 1 | 0.01987 | BP | Amillepora00093-RA | 0.220993171 |
| GO:0048268 | clathrin coat assembly | 1 | 1 | 0.01987 | BP | Amillepora20340-RA | 0.220993171 |
| GO:0060384 | innervation | 1 | 1 | 0.01987 | BP | Amillepora08533-RA | 0.220993171 |
| GO:0046604 | positive regulation of mitotic centrosome separation | 1 | 1 | 0.01987 | BP | Amillepora22273-RA | 0.220993171 |
| GO:0006382 | adenosine to inosine editing | 1 | 1 | 0.01987 | BP | Amillepora08533-RA | 0.220993171 |
| GO:0032461 | positive regulation of protein oligomerization | 1 | 1 | 0.01987 | BP | Amillepora14835-RA | 0.220993171 |
| GO:0032462 | regulation of protein homooligomerization | 1 | 1 | 0.01987 | BP | Amillepora14835-RA | 0.220993171 |
| GO:0032464 | positive regulation of protein homooligomerization | 1 | 1 | 0.01987 | BP | Amillepora14835-RA | 0.220993171 |
| GO:0046440 | L-lysine metabolic process | 1 | 1 | 0.01987 | BP | Amillepora16854-RA | 0.220993171 |
| GO:0032459 | regulation of protein oligomerization | 1 | 1 | 0.01987 | BP | Amillepora14835-RA | 0.220993171 |
| GO:0051897 | positive regulation of protein kinase B signaling | 1 | 1 | 0.01987 | BP | Amillepora03480-RA | 0.220993171 |
| GO:1903391 | regulation of adherens junction organization | 1 | 1 | 0.01987 | BP | Amillepora21104-RA | 0.220993171 |
| GO:1903392 | negative regulation of adherens junction organization | 1 | 1 | 0.01987 | BP | Amillepora21104-RA | 0.220993171 |
| GO:0048764 | trichoblast maturation | 1 | 1 | 0.01987 | BP | Amillepora22668-RA | 0.220993171 |
| GO:0048765 | root hair cell differentiation | 1 | 1 | 0.01987 | BP | Amillepora22668-RA | 0.220993171 |
| GO:0048767 | root hair elongation | 1 | 1 | 0.01987 | BP | Amillepora22668-RA | 0.220993171 |
| GO:0019878 | lysine biosynthetic process via aminoadipic acid | 1 | 1 | 0.01987 | BP | Amillepora16854-RA | 0.220993171 |
| GO:0034087 | establishment of mitotic sister chromatid cohesion | 1 | 1 | 0.01987 | BP | Amillepora21960-RA | 0.220993171 |
| GO:0030194 | positive regulation of blood coagulation | 1 | 1 | 0.01987 | BP | Amillepora03480-RA | 0.220993171 |
| GO:0033512 | L-lysine catabolic process to acetyl-CoA via saccharopine | 1 | 1 | 0.01987 | BP | Amillepora16854-RA | 0.220993171 |
| GO:0021561 | facial nerve development | 1 | 1 | 0.01987 | BP | Amillepora08533-RA | 0.220993171 |
| GO:0032197 | transposition, RNA-mediated | 1 | 1 | 0.01987 | BP | Amillepora18353-RA | 0.220993171 |
| GO:0021566 | hypoglossal nerve development | 1 | 1 | 0.01987 | BP | Amillepora08533-RA | 0.220993171 |
| GO:0090713 | immunological memory process | 1 | 1 | 0.01987 | BP | Amillepora10499-RA | 0.220993171 |
| GO:0090715 | immunological memory formation process | 1 | 1 | 0.01987 | BP | Amillepora10499-RA | 0.220993171 |
| GO:0034729 | histone H3-K79 methylation | 1 | 1 | 0.01987 | BP | Amillepora25070-RA | 0.220993171 |
| GO:0006438 | valyl-tRNA aminoacylation | 1 | 1 | 0.01987 | BP | Amillepora01328-RA | 0.220993171 |
| GO:0015916 | fatty-acyl-CoA transport | 1 | 1 | 0.01987 | BP | Amillepora19435-RA | 0.220993171 |
| GO:0097049 | motor neuron apoptotic process | 1 | 1 | 0.01987 | BP | Amillepora08533-RA | 0.220993171 |
| GO:1902992 | negative regulation of amyloid precursor protein catabolic process | 1 | 1 | 0.01987 | BP | Amillepora15763-RA | 0.220993171 |
| GO:1902994 | regulation of phospholipid efflux | 1 | 1 | 0.01987 | BP | Amillepora14835-RA | 0.220993171 |
| GO:1902995 | positive regulation of phospholipid efflux | 1 | 1 | 0.01987 | BP | Amillepora14835-RA | 0.220993171 |
| GO:0033690 | positive regulation of osteoblast proliferation | 1 | 1 | 0.01987 | BP | Amillepora03480-RA | 0.220993171 |
| GO:0010573 | vascular endothelial growth factor production | 1 | 1 | 0.01987 | BP | Amillepora03480-RA | 0.220993171 |
| GO:0010574 | regulation of vascular endothelial growth factor production | 1 | 1 | 0.01987 | BP | Amillepora03480-RA | 0.220993171 |
| GO:0010575 | positive regulation of vascular endothelial growth factor production | 1 | 1 | 0.01987 | BP | Amillepora03480-RA | 0.220993171 |
| GO:0010525 | regulation of transposition, RNA-mediated | 1 | 1 | 0.01987 | BP | Amillepora18353-RA | 0.220993171 |
| GO:0010526 | negative regulation of transposition, RNA-mediated | 1 | 1 | 0.01987 | BP | Amillepora18353-RA | 0.220993171 |
| GO:0061780 | mitotic cohesin loading | 1 | 1 | 0.01987 | BP | Amillepora21960-RA | 0.220993171 |
| GO:1902960 | negative regulation of aspartic-type endopeptidase activity involved in amyloid precursor protein catabolic proces | 1 | 1 | 0.01987 | BP | Amillepora15763-RA | 0.220993171 |
| GO:0061028 | establishment of endothelial barrier | 1 | 1 | 0.01987 | BP | Amillepora03480-RA | 0.220993171 |
| GO:1902959 | regulation of aspartic-type endopeptidase activity involved in amyloid precursor protein catabolic process | 1 | 1 | 0.01987 | BP | Amillepora15763-RA | 0.220993171 |
| GO:0002501 | peptide antigen assembly with MHC protein complex | 1 | 1 | 0.01987 | BP | Amillepora00093-RA | 0.220993171 |
| GO:0002502 | peptide antigen assembly with MHC class I protein complex | 1 | 1 | 0.01987 | BP | Amillepora00093-RA | 0.220993171 |
| GO:0002396 | MHC protein complex assembly | 1 | 1 | 0.01987 | BP | Amillepora00093-RA | 0.220993171 |
| GO:0002397 | MHC class I protein complex assembly | 1 | 1 | 0.01987 | BP | Amillepora00093-RA | 0.220993171 |
| GO:0033688 | regulation of osteoblast proliferation | 1 | 1 | 0.01987 | BP | Amillepora03480-RA | 0.220993171 |
| GO:0001174 | transcriptional start site selection at RNA polymerase II promoter | 1 | 1 | 0.01987 | BP | Amillepora24928-RA | 0.220993171 |
| GO:0001178 | regulation of transcriptional start site selection at RNA polymerase II promoter | 1 | 1 | 0.01987 | BP | Amillepora24928-RA | 0.220993171 |
| GO:0045875 | negative regulation of sister chromatid cohesion | 1 | 1 | 0.01987 | BP | Amillepora21960-RA | 0.220993171 |
| GO:0001894 | tissue homeostasis | 30 | 3 | 0.02108 | BP | Amillepora06397-RA, Amil | 0.233029818 |
| GO:0051248 | negative regulation of protein metabolic process | 118 | 8 | 0.02163 | BP | Amillepora11097-RA, Amil | 0.237669398 |
| GO:0034976 | response to endoplasmic reticulum stress | 54 | 4 | 0.02196 | BP | Amillepora11097-RA, Amil | 0.238453156 |
| GO:0030029 | actin filament-based process | 110 | 6 | 0.02198 | BP | Amillepora11097-RA, Amil | 0.238453156 |
| GO:0001895 | retina homeostasis | 12 | 2 | 0.02272 | BP | Amillepora00666-RA, Amil | 0.238453156 |
| GO:0030855 | epithelial cell differentiation | 58 | 6 | 0.02369 | BP | Amillepora21104-RA, Amil | 0.238453156 |
| GO:0051172 | negative regulation of nitrogen compound metabolic process | 256 | 12 | 0.02429 | BP | Amillepora23389-RA, Amil | 0.238453156 |
| GO:0006457 | protein folding | 83 | 5 | 0.02435 | BP | Amillepora26017-RA, Amil | 0.238453156 |
| GO:0048731 | system development | 654 | 23 | 0.02439 | BP | Amillepora15159-RA, Amil | 0.238453156 |
| GO:0006355 | regulation of DNA-templated transcription | 635 | 20 | 0.02572 | BP | Amillepora24928-RA, Amil | 0.238453156 |
| GO:1903506 | regulation of nucleic acid-templated transcription | 636 | 20 | 0.02611 | BP | Amillepora24928-RA, Amil | 0.238453156 |
| GO:0051168 | nuclear export | 13 | 2 | 0.0265 | BP | Amillepora24928-RA, Amil | 0.238453156 |
| GO:0035270 | endocrine system development | 13 | 2 | 0.0265 | BP | Amillepora05687-RA, Amil | 0.238453156 |
| GO:2001141 | regulation of RNA biosynthetic process | 638 | 20 | 0.02689 | BP | Amillepora24928-RA, Amil | 0.238453156 |
| GO:0036503 | ERAD pathway | 33 | 3 | 0.02714 | BP | Amillepora07010-RA, Amil | 0.238453156 |
| GO:0010556 | regulation of macromolecule biosynthetic process | 723 | 22 | 0.02778 | BP | Amillepora17925-RA, Amil | 0.238453156 |
| GO:0032880 | regulation of protein localization | 86 | 5 | 0.02787 | BP | Amillepora10499-RA, Amil | 0.238453156 |
| GO:0010605 | negative regulation of macromolecule metabolic process | 338 | 14 | 0.02898 | BP | Amillepora05037-RA, Amil | 0.238453156 |
| GO:0140014 | mitotic nuclear division | 59 | 4 | 0.0293 | BP | Amillepora21960-RA, Amil | 0.238453156 |
| GO:0071396 | cellular response to lipid | 34 | 3 | 0.02935 | BP | Amillepora21104-RA, Amil | 0.238453156 |
| GO:0065009 | regulation of molecular function | 417 | 16 | 0.02991 | BP | Amillepora05037-RA, Amil | 0.238453156 |
| GO:0071495 | cellular response to endogenous stimulus | 151 | 7 | 0.03051 | BP | Amillepora00093-RA, Amil | 0.238453156 |
| GO:0006900 | vesicle budding from membrane | 14 | 2 | 0.03052 | BP | Amillepora15763-RA, Amil | 0.238453156 |
| GO:0098542 | defense response to other organism | 119 | 6 | 0.03085 | BP | Amillepora15159-RA, Amil | 0.238453156 |
| GO:0006955 | immune response | 152 | 7 | 0.03147 | BP | Amillepora22025-RA, Amil | 0.238453156 |
| GO:0033365 | protein localization to organelle | 161 | 9 | 0.03151 | BP | Amillepora19361-RA, Amil | 0.238453156 |
| GO:2000113 | negative regulation of cellular macromolecule biosynthetic process | 120 | 6 | 0.03197 | BP | Amillepora11097-RA, Amil | 0.238453156 |
| GO:0042221 | response to chemical | 542 | 19 | 0.03231 | BP | Amillepora11351-RA, Amil | 0.238453156 |
| GO:0006790 | sulfur compound metabolic process | 153 | 7 | 0.03245 | BP | Amillepora16446-RA, Amil | 0.238453156 |
| GO:0006351 | DNA-templated transcription | 694 | 21 | 0.03305 | BP | Amillepora15159-RA, Amil | 0.238453156 |

| | | | | | | | |
|---|---|---|---|---|---|---|---|
| GO:0006749 | glutathione metabolic process | 36 | 3 | 0.03403 | BP | Amillepora04345-RA, Amil | 0.238453156 |
| GO:0033993 | response to lipid | 62 | 4 | 0.03433 | BP | Amillepora21104-RA, Amil | 0.238453156 |
| GO:0009888 | tissue development | 244 | 15 | 0.03468 | BP | Amillepora16669-RA, Amil | 0.238453156 |
| GO:0055001 | muscle cell development | 15 | 2 | 0.03476 | BP | Amillepora11097-RA, Amil | 0.238453156 |
| GO:0051289 | protein homotetramerization | 15 | 2 | 0.03476 | BP | Amillepora22668-RA, Amil | 0.238453156 |
| GO:0046364 | monosaccharide biosynthetic process | 15 | 2 | 0.03476 | BP | Amillepora10499-RA, Amil | 0.238453156 |
| GO:0007062 | sister chromatid cohesion | 15 | 2 | 0.03476 | BP | Amillepora08867-RA, Amil | 0.238453156 |
| GO:0005977 | glycogen metabolic process | 15 | 2 | 0.03476 | BP | Amillepora16446-RA, Amil | 0.238453156 |
| GO:0019827 | stem cell population maintenance | 15 | 2 | 0.03476 | BP | Amillepora16298-RA, Amil | 0.238453156 |
| GO:0006112 | energy reserve metabolic process | 15 | 2 | 0.03476 | BP | Amillepora16446-RA, Amil | 0.238453156 |
| GO:1902531 | regulation of intracellular signal transduction | 156 | 7 | 0.03551 | BP | Amillepora22025-RA, Amil | 0.238453156 |
| GO:0048285 | organelle fission | 92 | 5 | 0.03585 | BP | Amillepora03553-RA, Amil | 0.238453156 |
| GO:0031326 | regulation of cellular biosynthetic process | 743 | 22 | 0.03626 | BP | Amillepora17925-RA, Amil | 0.238453156 |
| GO:0097659 | nucleic acid-templated transcription | 704 | 21 | 0.03778 | BP | Amillepora15159-RA, Amil | 0.238453156 |
| GO:1902679 | negative regulation of RNA biosynthetic process | 125 | 6 | 0.03791 | BP | Amillepora11097-RA, Amil | 0.238453156 |
| GO:1903507 | negative regulation of nucleic acid-templated transcription | 125 | 6 | 0.03791 | BP | Amillepora11097-RA, Amil | 0.238453156 |
| GO:0051656 | establishment of organelle localization | 64 | 4 | 0.03795 | BP | Amillepora15763-RA, Amil | 0.238453156 |
| GO:2000112 | regulation of cellular macromolecule biosynthetic process | 705 | 21 | 0.03827 | BP | Amillepora18353-RA, Amil | 0.238453156 |
| GO:0010558 | negative regulation of macromolecule biosynthetic process | 159 | 7 | 0.03876 | BP | Amillepora11097-RA, Amil | 0.238453156 |
| GO:0009615 | response to virus | 38 | 3 | 0.03908 | BP | Amillepora04345-RA, Amil | 0.238453156 |
| GO:0043434 | response to peptide hormone | 38 | 3 | 0.03908 | BP | Amillepora02951-RA, Amil | 0.238453156 |
| GO:0035303 | regulation of dephosphorylation | 16 | 2 | 0.03922 | BP | Amillepora21990-RA, Amil | 0.238453156 |
| GO:0007626 | locomotory behavior | 16 | 2 | 0.03922 | BP | Amillepora20112-RA, Amil | 0.238453156 |
| GO:0044042 | glucan metabolic process | 16 | 2 | 0.03922 | BP | Amillepora16446-RA, Amil | 0.238453156 |
| GO:0006909 | phagocytosis | 16 | 2 | 0.03922 | BP | Amillepora21104-RA, Amil | 0.238453156 |
| GO:0098727 | maintenance of cell number | 16 | 2 | 0.03922 | BP | Amillepora16298-RA, Amil | 0.238453156 |
| GO:0045664 | regulation of neuron differentiation | 16 | 2 | 0.03922 | BP | Amillepora21104-RA, Amil | 0.238453156 |
| GO:0006073 | cellular glucan metabolic process | 16 | 2 | 0.03922 | BP | Amillepora16446-RA, Amil | 0.238453156 |
| GO:0006116 | NADH oxidation | 2 | 1 | 0.03935 | BP | Amillepora10499-RA | 0.238453156 |
| GO:0016185 | synaptic vesicle budding from presynaptic endocytic zone membrane | 2 | 1 | 0.03935 | BP | Amillepora20340-RA | 0.238453156 |
| GO:0002688 | regulation of leukocyte chemotaxis | 2 | 1 | 0.03935 | BP | Amillepora00093-RA | 0.238453156 |
| GO:1900026 | positive regulation of substrate adhesion-dependent cell spreading | 2 | 1 | 0.03935 | BP | Amillepora00093-RA | 0.238453156 |
| GO:1900048 | positive regulation of hemostasis | 2 | 1 | 0.03935 | BP | Amillepora03480-RA | 0.238453156 |
| GO:0021602 | cranial nerve morphogenesis | 2 | 1 | 0.03935 | BP | Amillepora08533-RA | 0.238453156 |
| GO:0051674 | localization of cell | 2 | 1 | 0.03935 | BP | Amillepora11351-RA | 0.238453156 |
| GO:0030200 | heparan sulfate proteoglycan catabolic process | 2 | 1 | 0.03935 | BP | Amillepora03480-RA | 0.238453156 |
| GO:0050820 | positive regulation of coagulation | 2 | 1 | 0.03935 | BP | Amillepora03480-RA | 0.238453156 |
| GO:0060217 | hemangioblast cell differentiation | 2 | 1 | 0.03935 | BP | Amillepora05687-RA | 0.238453156 |
| GO:0045579 | positive regulation of B cell differentiation | 2 | 1 | 0.03935 | BP | Amillepora15111-RA | 0.238453156 |
| GO:0150172 | regulation of phosphatidylcholine metabolic process | 2 | 1 | 0.03935 | BP | Amillepora14835-RA | 0.238453156 |
| GO:0001946 | lymphangiogenesis | 2 | 1 | 0.03935 | BP | Amillepora05687-RA | 0.238453156 |
| GO:0033198 | response to ATP | 2 | 1 | 0.03935 | BP | Amillepora22291-RA | 0.238453156 |
| GO:0044703 | multi-organism reproductive process | 2 | 1 | 0.03935 | BP | Amillepora00093-RA | 0.238453156 |
| GO:0044706 | multi-multicellular organism process | 2 | 1 | 0.03935 | BP | Amillepora00093-RA | 0.238453156 |
| GO:0060055 | angiogenesis involved in wound healing | 2 | 1 | 0.03935 | BP | Amillepora03480-RA | 0.238453156 |
| GO:0002407 | dendritic cell chemotaxis | 2 | 1 | 0.03935 | BP | Amillepora00093-RA | 0.238453156 |
| GO:0048385 | regulation of retinoic acid receptor signaling pathway | 2 | 1 | 0.03935 | BP | Amillepora00093-RA | 0.238453156 |
| GO:0006086 | acetyl-CoA biosynthetic process from pyruvate | 2 | 1 | 0.03935 | BP | Amillepora03530-RA | 0.238453156 |
| GO:0070142 | synaptic vesicle budding | 2 | 1 | 0.03935 | BP | Amillepora20340-RA | 0.238453156 |
| GO:0055090 | acylglycerol homeostasis | 2 | 1 | 0.03935 | BP | Amillepora16446-RA | 0.238453156 |
| GO:0055091 | phospholipid homeostasis | 2 | 1 | 0.03935 | BP | Amillepora14835-RA | 0.238453156 |
| GO:0033144 | negative regulation of intracellular steroid hormone receptor signaling pathway | 2 | 1 | 0.03935 | BP | Amillepora00093-RA | 0.238453156 |
| GO:2000048 | negative regulation of cell-cell adhesion mediated by cadherin | 2 | 1 | 0.03935 | BP | Amillepora21104-RA | 0.238453156 |
| GO:2000001 | regulation of DNA damage checkpoint | 2 | 1 | 0.03935 | BP | Amillepora09972-RA | 0.238453156 |
| GO:0090303 | positive regulation of wound healing | 2 | 1 | 0.03935 | BP | Amillepora03480-RA | 0.238453156 |
| GO:0042989 | sequestering of actin monomers | 2 | 1 | 0.03935 | BP | Amillepora19558-RA | 0.238453156 |
| GO:0036303 | lymph vessel morphogenesis | 2 | 1 | 0.03935 | BP | Amillepora05687-RA | 0.238453156 |
| GO:1901163 | regulation of trophoblast cell migration | 2 | 1 | 0.03935 | BP | Amillepora00093-RA | 0.238453156 |
| GO:1901164 | negative regulation of trophoblast cell migration | 2 | 1 | 0.03935 | BP | Amillepora00093-RA | 0.238453156 |
| GO:0007274 | neuromuscular synaptic transmission | 2 | 1 | 0.03935 | BP | Amillepora08533-RA | 0.238453156 |
| GO:0042754 | negative regulation of circadian rhythm | 2 | 1 | 0.03935 | BP | Amillepora23389-RA | 0.238453156 |
| GO:0061450 | trophoblast cell migration | 2 | 1 | 0.03935 | BP | Amillepora00093-RA | 0.238453156 |
| GO:0030539 | male genitalia development | 2 | 1 | 0.03935 | BP | Amillepora21104-RA | 0.238453156 |
| GO:0099402 | plant organ development | 2 | 1 | 0.03935 | BP | Amillepora22668-RA | 0.238453156 |
| GO:0039529 | RIG-I signaling pathway | 2 | 1 | 0.03935 | BP | Amillepora22025-RA | 0.238453156 |
| GO:0007063 | regulation of sister chromatid cohesion | 2 | 1 | 0.03935 | BP | Amillepora21960-RA | 0.238453156 |
| GO:0039535 | regulation of RIG-I signaling pathway | 2 | 1 | 0.03935 | BP | Amillepora22025-RA | 0.238453156 |
| GO:0039536 | negative regulation of RIG-I signaling pathway | 2 | 1 | 0.03935 | BP | Amillepora22025-RA | 0.238453156 |
| GO:0046327 | glycerol biosynthetic process from pyruvate | 2 | 1 | 0.03935 | BP | Amillepora10499-RA | 0.238453156 |
| GO:0016553 | base conversion or substitution editing | 2 | 1 | 0.03935 | BP | Amillepora08533-RA | 0.238453156 |
| GO:0051797 | regulation of hair follicle development | 2 | 1 | 0.03935 | BP | Amillepora03480-RA | 0.238453156 |
| GO:0060390 | regulation of SMAD protein signal transduction | 2 | 1 | 0.03935 | BP | Amillepora21104-RA | 0.238453156 |
| GO:0060393 | regulation of pathway-restricted SMAD protein phosphorylation | 2 | 1 | 0.03935 | BP | Amillepora21104-RA | 0.238453156 |
| GO:0046618 | xenobiotic export from cell | 2 | 1 | 0.03935 | BP | Amillepora14835-RA | 0.238453156 |
| GO:0061158 | 3'-UTR-mediated mRNA destabilization | 2 | 1 | 0.03935 | BP | Amillepora18353-RA | 0.238453156 |
| GO:0010039 | response to iron ion | 2 | 1 | 0.03935 | BP | Amillepora21104-RA | 0.238453156 |
| GO:0071675 | regulation of mononuclear cell migration | 2 | 1 | 0.03935 | BP | Amillepora00093-RA | 0.238453156 |
| GO:0071677 | positive regulation of mononuclear cell migration | 2 | 1 | 0.03935 | BP | Amillepora00093-RA | 0.238453156 |
| GO:0060389 | pathway-restricted SMAD protein phosphorylation | 2 | 1 | 0.03935 | BP | Amillepora21104-RA | 0.238453156 |
| GO:1903566 | positive regulation of protein localization to cilium | 2 | 1 | 0.03935 | BP | Amillepora00666-RA | 0.238453156 |
| GO:0070841 | inclusion body assembly | 2 | 1 | 0.03935 | BP | Amillepora11097-RA | 0.238453156 |
| GO:0017003 | protein-heme linkage | 2 | 1 | 0.03935 | BP | Amillepora25339-RA | 0.238453156 |
| GO:0017006 | protein-tetrapyrrole linkage | 2 | 1 | 0.03935 | BP | Amillepora25339-RA | 0.238453156 |
| GO:2001140 | positive regulation of phospholipid transport | 2 | 1 | 0.03935 | BP | Amillepora14835-RA | 0.238453156 |
| GO:2001138 | regulation of phospholipid transport | 2 | 1 | 0.03935 | BP | Amillepora14835-RA | 0.238453156 |
| GO:0010862 | positive regulation of pathway-restricted SMAD protein phosphorylation | 2 | 1 | 0.03935 | BP | Amillepora21104-RA | 0.238453156 |
| GO:0010875 | positive regulation of cholesterol efflux | 2 | 1 | 0.03935 | BP | Amillepora14835-RA | 0.238453156 |
| GO:1905392 | plant organ morphogenesis | 2 | 1 | 0.03935 | BP | Amillepora22668-RA | 0.238453156 |
| GO:0090083 | regulation of inclusion body assembly | 2 | 1 | 0.03935 | BP | Amillepora11097-RA | 0.238453156 |
| GO:0090084 | negative regulation of inclusion body assembly | 2 | 1 | 0.03935 | BP | Amillepora11097-RA | 0.238453156 |
| GO:0034085 | establishment of sister chromatid cohesion | 2 | 1 | 0.03935 | BP | Amillepora21960-RA | 0.238453156 |
| GO:0030167 | proteoglycan catabolic process | 2 | 1 | 0.03935 | BP | Amillepora03480-RA | 0.238453156 |
| GO:0007565 | female pregnancy | 2 | 1 | 0.03935 | BP | Amillepora00093-RA | 0.238453156 |
| GO:0007566 | embryo implantation | 2 | 1 | 0.03935 | BP | Amillepora00093-RA | 0.238453156 |
| GO:0006437 | tyrosyl-tRNA aminoacylation | 2 | 1 | 0.03935 | BP | Amillepora20065-RA | 0.238453156 |
| GO:0009561 | megagametogenesis | 2 | 1 | 0.03935 | BP | Amillepora20323-RA | 0.238453156 |
| GO:0009553 | embryo sac development | 2 | 1 | 0.03935 | BP | Amillepora20323-RA | 0.238453156 |
| GO:0006405 | RNA export from nucleus | 2 | 1 | 0.03935 | BP | Amillepora24928-RA | 0.238453156 |

| | | | | | | | |
|---|---|---|---|---|---|---|---|
| GO:1905634 | regulation of protein localization to chromatin | 2 | 1 | 0.03935 | BP | Amillepora21960-RA | 0.238453156 |
| GO:0010528 | regulation of transposition | 2 | 1 | 0.03935 | BP | Amillepora18353-RA | 0.238453156 |
| GO:0010529 | negative regulation of transposition | 2 | 1 | 0.03935 | BP | Amillepora18353-RA | 0.238453156 |
| GO:0018063 | cytochrome c-heme linkage | 2 | 1 | 0.03935 | BP | Amillepora25339-RA | 0.238453156 |
| GO:0070328 | triglyceride homeostasis | 2 | 1 | 0.03935 | BP | Amillepora16446-RA | 0.238453156 |
| GO:0048875 | chemical homeostasis within a tissue | 2 | 1 | 0.03935 | BP | Amillepora14835-RA | 0.238453156 |
| GO:0001173 | DNA-templated transcriptional start site selection | 2 | 1 | 0.03935 | BP | Amillepora24928-RA | 0.238453156 |
| GO:0002690 | positive regulation of leukocyte chemotaxis | 2 | 1 | 0.03935 | BP | Amillepora00093-RA | 0.238453156 |
| GO:0019543 | propionate catabolic process | 2 | 1 | 0.03935 | BP | Amillepora10499-RA | 0.238453156 |
| GO:0016070 | RNA metabolic process | 1130 | 33 | 0.04107 | BP | Amillepora09499-RA, Amil | 0.245609585 |
| GO:0051648 | vesicle localization | 39 | 3 | 0.04173 | BP | Amillepora20340-RA, Amil | 0.245609585 |
| GO:0060249 | anatomical structure homeostasis | 39 | 3 | 0.04173 | BP | Amillepora06397-RA, Amil | 0.245609585 |
| GO:0034754 | cellular hormone metabolic process | 39 | 3 | 0.04173 | BP | Amillepora05711-RA, Amil | 0.245609585 |
| GO:0032103 | positive regulation of response to external stimulus | 39 | 3 | 0.04173 | BP | Amillepora15159-RA, Amil | 0.245609585 |
| GO:0032101 | regulation of response to external stimulus | 96 | 5 | 0.04188 | BP | Amillepora00093-RA, Amil | 0.245609585 |
| GO:0030036 | actin cytoskeleton organization | 97 | 5 | 0.04348 | BP | Amillepora11097-RA, Amil | 0.245609585 |
| GO:0006575 | cellular modified amino acid metabolic process | 97 | 5 | 0.04348 | BP | Amillepora04345-RA, Amil | 0.245609585 |
| GO:0007059 | chromosome segregation | 67 | 4 | 0.04376 | BP | Amillepora12757-RA, Amil | 0.245609585 |
| GO:1903037 | regulation of leukocyte cell-cell adhesion | 17 | 2 | 0.04388 | BP | Amillepora10499-RA, Amil | 0.245609585 |
| GO:0051262 | protein tetramerization | 17 | 2 | 0.04388 | BP | Amillepora22668-RA, Amil | 0.245609585 |
| GO:0022409 | positive regulation of cell-cell adhesion | 17 | 2 | 0.04388 | BP | Amillepora10499-RA, Amil | 0.245609585 |
| GO:0097722 | sperm motility | 17 | 2 | 0.04388 | BP | Amillepora11351-RA, Amil | 0.245609585 |
| GO:0009719 | response to endogenous stimulus | 164 | 7 | 0.0446 | BP | Amillepora00093-RA, Amil | 0.245609585 |
| GO:0032774 | RNA biosynthetic process | 717 | 21 | 0.04464 | BP | Amillepora15159-RA, Amil | 0.245609585 |
| GO:0050790 | regulation of catalytic activity | 361 | 14 | 0.04492 | BP | Amillepora05037-RA, Amil | 0.245609585 |
| GO:0009605 | response to external stimulus | 292 | 17 | 0.04504 | BP | Amillepora11351-RA, Amil | 0.245609585 |
| GO:0007169 | transmembrane receptor protein tyrosine kinase signaling pathway | 68 | 4 | 0.04581 | BP | Amillepora18085-RA, Amil | 0.245609585 |
| GO:0032870 | cellular response to hormone stimulus | 68 | 4 | 0.04581 | BP | Amillepora02951-RA, Amil | 0.245609585 |
| GO:0009889 | regulation of biosynthetic process | 752 | 24 | 0.04613 | BP | Amillepora17925-RA, Amil | 0.245609585 |
| GO:0060341 | regulation of cellular localization | 99 | 5 | 0.04678 | BP | Amillepora10499-RA, Amil | 0.245609585 |
| GO:1901652 | response to peptide | 41 | 3 | 0.04732 | BP | Amillepora02951-RA, Amil | 0.245609585 |
| GO:0060041 | retina development in camera-type eye | 41 | 3 | 0.04732 | BP | Amillepora15159-RA, Amil | 0.245609585 |
| GO:1903047 | mitotic cell cycle process | 132 | 6 | 0.04737 | BP | Amillepora11105-RA, Amil | 0.245609585 |
| GO:0018108 | peptidyl-tyrosine phosphorylation | 69 | 4 | 0.0479 | BP | Amillepora21104-RA, Amil | 0.245609585 |
| GO:0009892 | negative regulation of metabolic process | 365 | 14 | 0.04821 | BP | Amillepora05037-RA, Amil | 0.245609585 |
| GO:0031324 | negative regulation of cellular metabolic process | 239 | 9 | 0.04822 | BP | Amillepora23389-RA, Amil | 0.245609585 |
| GO:0097435 | supramolecular fiber organization | 100 | 5 | 0.04849 | BP | Amillepora11097-RA, Amil | 0.245609585 |
| GO:0071773 | cellular response to BMP stimulus | 18 | 2 | 0.04874 | BP | Amillepora21104-RA, Amil | 0.245609585 |
| GO:0002064 | epithelial cell development | 18 | 2 | 0.04874 | BP | Amillepora21104-RA, Amil | 0.245609585 |
| GO:0032606 | type I interferon production | 18 | 2 | 0.04874 | BP | Amillepora22025-RA, Amil | 0.245609585 |
| GO:1903532 | positive regulation of secretion by cell | 18 | 2 | 0.04874 | BP | Amillepora15763-RA, Amil | 0.245609585 |
| GO:0032479 | regulation of type I interferon production | 18 | 2 | 0.04874 | BP | Amillepora22025-RA, Amil | 0.245609585 |
| GO:0046173 | polyol biosynthetic process | 18 | 2 | 0.04874 | BP | Amillepora16062-RA, Amil | 0.245609585 |
| GO:0006357 | regulation of transcription by RNA polymerase II | 316 | 11 | 0.0495 | BP | Amillepora10714-RA, Amil | 0.245609585 |
| GO:0001654 | eye development | 70 | 4 | 0.05005 | BP | Amillepora21104-RA, Amil | 0.245609585 |
| GO:0040007 | growth | 101 | 5 | 0.05023 | BP | Amillepora22668-RA, Amil | 0.245609585 |
| GO:0006954 | inflammatory response | 42 | 3 | 0.05024 | BP | Amillepora04708-RA, Amil | 0.245609585 |
| GO:0031667 | response to nutrient levels | 42 | 3 | 0.05024 | BP | Amillepora16446-RA, Amil | 0.245609585 |
| GO:0030182 | neuron differentiation | 169 | 7 | 0.05098 | BP | Amillepora00093-RA, Amil | 0.245609585 |
| GO:0051253 | negative regulation of RNA metabolic process | 135 | 6 | 0.05183 | BP | Amillepora11097-RA, Amil | 0.245609585 |
| GO:0032501 | multicellular organismal process | 985 | 35 | 0.05199 | BP | Amillepora17925-RA, Amil | 0.245609585 |
| GO:0031327 | negative regulation of cellular biosynthetic process | 171 | 7 | 0.05369 | BP | Amillepora05687-RA, Amil | 0.245609585 |
| GO:0050708 | regulation of protein secretion | 19 | 2 | 0.05378 | BP | Amillepora15763-RA, Amil | 0.245609585 |
| GO:0030217 | T cell differentiation | 19 | 2 | 0.05378 | BP | Amillepora10499-RA, Amil | 0.245609585 |
| GO:0050863 | regulation of T cell activation | 19 | 2 | 0.05378 | BP | Amillepora10499-RA, Amil | 0.245609585 |
| GO:0006090 | pyruvate metabolic process | 19 | 2 | 0.05378 | BP | Amillepora10499-RA, Amil | 0.245609585 |
| GO:0071772 | response to BMP | 19 | 2 | 0.05378 | BP | Amillepora21104-RA, Amil | 0.245609585 |
| GO:0007159 | leukocyte cell-cell adhesion | 19 | 2 | 0.05378 | BP | Amillepora10499-RA, Amil | 0.245609585 |
| GO:0050657 | nucleic acid transport | 19 | 2 | 0.05378 | BP | Amillepora24928-RA, Amil | 0.245609585 |
| GO:0050658 | RNA transport | 19 | 2 | 0.05378 | BP | Amillepora24928-RA, Amil | 0.245609585 |
| GO:0051236 | establishment of RNA localization | 19 | 2 | 0.05378 | BP | Amillepora24928-RA, Amil | 0.245609585 |
| GO:0051047 | positive regulation of secretion | 19 | 2 | 0.05378 | BP | Amillepora15763-RA, Amil | 0.245609585 |
| GO:0051129 | negative regulation of cellular component organization | 72 | 4 | 0.05451 | BP | Amillepora11097-RA, Amil | 0.245609585 |
| GO:0150063 | visual system development | 72 | 4 | 0.05451 | BP | Amillepora15159-RA, Amil | 0.245609585 |
| GO:0000226 | microtubule cytoskeleton organization | 137 | 6 | 0.05494 | BP | Amillepora11105-RA, Amil | 0.245609585 |
| GO:0030203 | glycosaminoglycan metabolic process | 44 | 3 | 0.05634 | BP | Amillepora03480-RA, Amil | 0.245609585 |
| GO:0015931 | nucleobase-containing compound transport | 44 | 3 | 0.05634 | BP | Amillepora22273-RA, Amil | 0.245609585 |
| GO:0051174 | regulation of phosphorus metabolic process | 138 | 6 | 0.05654 | BP | Amillepora14835-RA, Amil | 0.245609585 |
| GO:0019220 | regulation of phosphate metabolic process | 138 | 6 | 0.05654 | BP | Amillepora14835-RA, Amil | 0.245609585 |
| GO:0016311 | dephosphorylation | 138 | 6 | 0.05654 | BP | Amillepora24931-RA, Amil | 0.245609585 |
| GO:0051234 | establishment of localization | 1380 | 40 | 0.05675 | BP | Amillepora22025-RA, Amil | 0.245609585 |
| GO:0051050 | positive regulation of transport | 73 | 4 | 0.05682 | BP | Amillepora10499-RA, Amil | 0.245609585 |
| GO:0045087 | innate immune response | 73 | 4 | 0.05682 | BP | Amillepora22668-RA, Amil | 0.245609585 |
| GO:0007017 | microtubule-based process | 247 | 9 | 0.05715 | BP | Amillepora05723-RA, Amil | 0.245609585 |
| GO:0006325 | chromatin organization | 105 | 5 | 0.05757 | BP | Amillepora22378-RA, Amil | 0.245609585 |
| GO:0015031 | protein transport | 364 | 12 | 0.05785 | BP | Amillepora23397-RA, Amil | 0.245609585 |
| GO:0002687 | positive regulation of leukocyte migration | 3 | 1 | 0.05844 | BP | Amillepora00093-RA | 0.245609585 |
| GO:0048806 | genitalia development | 3 | 1 | 0.05844 | BP | Amillepora21104-RA | 0.245609585 |
| GO:0042420 | dopamine catabolic process | 3 | 1 | 0.05844 | BP | Amillepora12729-RA | 0.245609585 |
| GO:0042424 | catecholamine catabolic process | 3 | 1 | 0.05844 | BP | Amillepora12729-RA | 0.245609585 |
| GO:1900024 | regulation of substrate adhesion-dependent cell spreading | 3 | 1 | 0.05844 | BP | Amillepora00093-RA | 0.245609585 |
| GO:0032958 | inositol phosphate biosynthetic process | 3 | 1 | 0.05844 | BP | Amillepora16062-RA | 0.245609585 |
| GO:0042417 | dopamine metabolic process | 3 | 1 | 0.05844 | BP | Amillepora12729-RA | 0.245609585 |
| GO:0048524 | positive regulation of viral process | 3 | 1 | 0.05844 | BP | Amillepora08533-RA | 0.245609585 |
| GO:0019626 | short-chain fatty acid catabolic process | 3 | 1 | 0.05844 | BP | Amillepora10499-RA | 0.245609585 |
| GO:0045577 | regulation of B cell differentiation | 3 | 1 | 0.05844 | BP | Amillepora15111-RA | 0.245609585 |
| GO:0019614 | catechol-containing compound catabolic process | 3 | 1 | 0.05844 | BP | Amillepora12729-RA | 0.245609585 |
| GO:0032053 | ciliary basal body organization | 3 | 1 | 0.05844 | BP | Amillepora00666-RA | 0.245609585 |
| GO:0001945 | lymph vessel development | 3 | 1 | 0.05844 | BP | Amillepora05687-RA | 0.245609585 |
| GO:0062207 | regulation of pattern recognition receptor signaling pathway | 3 | 1 | 0.05844 | BP | Amillepora22025-RA | 0.245609585 |
| GO:1901317 | regulation of flagellated sperm motility | 3 | 1 | 0.05844 | BP | Amillepora11351-RA | 0.245609585 |
| GO:0010447 | response to acidic pH | 3 | 1 | 0.05844 | BP | Amillepora10499-RA | 0.245609585 |
| GO:0048384 | retinoic acid receptor signaling pathway | 3 | 1 | 0.05844 | BP | Amillepora00093-RA | 0.245609585 |
| GO:0045669 | positive regulation of osteoblast differentiation | 3 | 1 | 0.05844 | BP | Amillepora21104-RA | 0.245609585 |
| GO:0009409 | response to cold | 3 | 1 | 0.05844 | BP | Amillepora04680-RA | 0.245609585 |
| GO:0031664 | regulation of lipopolysaccharide-mediated signaling pathway | 3 | 1 | 0.05844 | BP | Amillepora21104-RA | 0.245609585 |
| GO:0031666 | positive regulation of lipopolysaccharide-mediated signaling pathway | 3 | 1 | 0.05844 | BP | Amillepora21104-RA | 0.245609585 |
| GO:0019401 | alditol biosynthetic process | 3 | 1 | 0.05844 | BP | Amillepora10499-RA | 0.245609585 |

| | | | | | | | |
|---|---|---|---|---|---|---|---|
| GO:0034341 | response to interferon-gamma | 3 | 1 | 0.05844 | BP | Amillepora22668-RA | 0.245609585 |
| GO:0006734 | NADH metabolic process | 3 | 1 | 0.05844 | BP | Amillepora10499-RA | 0.245609585 |
| GO:0036336 | dendritic cell migration | 3 | 1 | 0.05844 | BP | Amillepora00093-RA | 0.245609585 |
| GO:0006700 | C21-steroid hormone biosynthetic process | 3 | 1 | 0.05844 | BP | Amillepora21104-RA | 0.245609585 |
| GO:0006705 | mineralocorticoid biosynthetic process | 3 | 1 | 0.05844 | BP | Amillepora21104-RA | 0.245609585 |
| GO:0003323 | type B pancreatic cell development | 3 | 1 | 0.05844 | BP | Amillepora21104-RA | 0.245609585 |
| GO:0055008 | cardiac muscle tissue morphogenesis | 3 | 1 | 0.05844 | BP | Amillepora16669-RA | 0.245609585 |
| GO:0003309 | type B pancreatic cell differentiation | 3 | 1 | 0.05844 | BP | Amillepora21104-RA | 0.245609585 |
| GO:0021952 | central nervous system projection neuron axonogenesis | 3 | 1 | 0.05844 | BP | Amillepora08533-RA | 0.245609585 |
| GO:0014823 | response to activity | 3 | 1 | 0.05844 | BP | Amillepora04680-RA | 0.245609585 |
| GO:0080182 | histone H3-K4 trimethylation | 3 | 1 | 0.05844 | BP | Amillepora16298-RA | 0.245609585 |
| GO:0039528 | cytoplasmic pattern recognition receptor signaling pathway in response to virus | 3 | 1 | 0.05844 | BP | Amillepora22025-RA | 0.245609585 |
| GO:0039531 | regulation of viral-induced cytoplasmic pattern recognition receptor signaling pathway | 3 | 1 | 0.05844 | BP | Amillepora22025-RA | 0.245609585 |
| GO:0039532 | negative regulation of viral-induced cytoplasmic pattern recognition receptor signaling pathway | 3 | 1 | 0.05844 | BP | Amillepora22025-RA | 0.245609585 |
| GO:0006589 | octopamine biosynthetic process | 3 | 1 | 0.05844 | BP | Amillepora12729-RA | 0.245609585 |
| GO:0009068 | aspartate family amino acid catabolic process | 3 | 1 | 0.05844 | BP | Amillepora16854-RA | 0.245609585 |
| GO:0046333 | octopamine metabolic process | 3 | 1 | 0.05844 | BP | Amillepora12729-RA | 0.245609585 |
| GO:1901224 | positive regulation of NIK/NF-kappaB signaling | 3 | 1 | 0.05844 | BP | Amillepora00093-RA | 0.245609585 |
| GO:0032341 | aldosterone metabolic process | 3 | 1 | 0.05844 | BP | Amillepora21104-RA | 0.245609585 |
| GO:0032342 | aldosterone biosynthetic process | 3 | 1 | 0.05844 | BP | Amillepora21104-RA | 0.245609585 |
| GO:0032344 | regulation of aldosterone metabolic process | 3 | 1 | 0.05844 | BP | Amillepora21104-RA | 0.245609585 |
| GO:0032346 | positive regulation of aldosterone metabolic process | 3 | 1 | 0.05844 | BP | Amillepora21104-RA | 0.245609585 |
| GO:0032347 | regulation of aldosterone biosynthetic process | 3 | 1 | 0.05844 | BP | Amillepora21104-RA | 0.245609585 |
| GO:0006691 | leukotriene metabolic process | 3 | 1 | 0.05844 | BP | Amillepora09624-RA | 0.245609585 |
| GO:0046602 | regulation of mitotic centrosome separation | 3 | 1 | 0.05844 | BP | Amillepora22273-RA | 0.245609585 |
| GO:0071346 | cellular response to interferon-gamma | 3 | 1 | 0.05844 | BP | Amillepora22668-RA | 0.245609585 |
| GO:0048916 | posterior lateral line development | 3 | 1 | 0.05844 | BP | Amillepora07010-RA | 0.245609585 |
| GO:0046471 | phosphatidylglycerol metabolic process | 3 | 1 | 0.05844 | BP | Amillepora14835-RA | 0.245609585 |
| GO:0051896 | regulation of protein kinase B signaling | 3 | 1 | 0.05844 | BP | Amillepora03480-RA | 0.245609585 |
| GO:0032480 | negative regulation of type I interferon production | 3 | 1 | 0.05844 | BP | Amillepora22025-RA | 0.245609585 |
| GO:0071468 | cellular response to acidic pH | 3 | 1 | 0.05844 | BP | Amillepora10499-RA | 0.245609585 |
| GO:0010874 | regulation of cholesterol efflux | 3 | 1 | 0.05844 | BP | Amillepora14835-RA | 0.245609585 |
| GO:0010811 | positive regulation of cell-substrate adhesion | 3 | 1 | 0.05844 | BP | Amillepora00093-RA | 0.245609585 |
| GO:0008212 | mineralocorticoid metabolic process | 3 | 1 | 0.05844 | BP | Amillepora21104-RA | 0.245609585 |
| GO:0045070 | positive regulation of viral genome replication | 3 | 1 | 0.05844 | BP | Amillepora08533-RA | 0.245609585 |
| GO:0071168 | protein localization to chromatin | 3 | 1 | 0.05844 | BP | Amillepora21960-RA | 0.245609585 |
| GO:0015939 | pantothenate metabolic process | 3 | 1 | 0.05844 | BP | Amillepora04708-RA | 0.245609585 |
| GO:0002544 | chronic inflammatory response | 3 | 1 | 0.05844 | BP | Amillepora04708-RA | 0.245609585 |
| GO:0070365 | hepatocyte differentiation | 3 | 1 | 0.05844 | BP | Amillepora10499-RA | 0.245609585 |
| GO:0015798 | myo-inositol transport | 3 | 1 | 0.05844 | BP | Amillepora18526-RA | 0.245609585 |
| GO:0050766 | positive regulation of phagocytosis | 3 | 1 | 0.05844 | BP | Amillepora00093-RA | 0.245609585 |
| GO:0042634 | regulation of hair cycle | 3 | 1 | 0.05844 | BP | Amillepora03480-RA | 0.245609585 |
| GO:0033687 | osteoblast proliferation | 3 | 1 | 0.05844 | BP | Amillepora03480-RA | 0.245609585 |
| GO:0006114 | glycerol biosynthetic process | 3 | 1 | 0.05844 | BP | Amillepora10499-RA | 0.245609585 |
| GO:0031589 | cell-substrate adhesion | 20 | 2 | 0.05899 | BP | Amillepora03480-RA, Amil | 0.24621913 |
| GO:2000241 | regulation of reproductive process | 20 | 2 | 0.05899 | BP | Amillepora11351-RA, Amil | 0.24621913 |
| GO:0044273 | sulfur compound catabolic process | 20 | 2 | 0.05899 | BP | Amillepora09624-RA, Amil | 0.24621913 |
| GO:0009890 | negative regulation of biosynthetic process | 175 | 7 | 0.05936 | BP | Amillepora05687-RA, Amil | 0.246820364 |
| GO:1903829 | positive regulation of protein localization | 45 | 3 | 0.05952 | BP | Amillepora15763-RA, Amil | 0.246820364 |
| GO:0071310 | cellular response to organic substance | 249 | 9 | 0.05954 | BP | Amillepora18085-RA, Amil | 0.246820364 |
| GO:0005515 | protein binding | 1339 | 47 | 0.000043 | MF | Amillepora17937-RA, Amil | 0.017458 |
| GO:1990269 | RNA polymerase II C-terminal domain phosphoserine binding | 2 | 2 | 0.00033 | MF | Amillepora16298-RA, Amil | 0.06699 |
| GO:0045504 | dynein heavy chain binding | 6 | 2 | 0.00475 | MF | Amillepora11351-RA, Amil | 0.225022424 |
| GO:0098772 | molecular function regulator activity | 316 | 13 | 0.00515 | MF | Amillepora27228-RA, Amil | 0.225022424 |
| GO:0008092 | cytoskeletal protein binding | 235 | 10 | 0.01091 | MF | Amillepora16717-RA, Amil | 0.225022424 |
| GO:0051082 | unfolded protein binding | 48 | 4 | 0.01122 | MF | Amillepora11097-RA, Amil | 0.225022424 |
| GO:0004869 | cysteine-type endopeptidase inhibitor activity | 11 | 2 | 0.01641 | MF | Amillepora09624-RA, Amil | 0.225022424 |
| GO:0003779 | actin binding | 113 | 6 | 0.01728 | MF | Amillepora22084-RA, Amil | 0.225022424 |
| GO:0008444 | CDP-diacylglycerol-glycerol-3-phosphate 3-phosphatidyltransferase activity | 1 | 1 | 0.01829 | MF | Amillepora15323-RA | 0.225022424 |
| GO:0071936 | coreceptor activity involved in Wnt signaling pathway | 1 | 1 | 0.01829 | MF | Amillepora13244-RA | 0.225022424 |
| GO:0004739 | pyruvate dehydrogenase (acetyl-transferring) activity | 1 | 1 | 0.01829 | MF | Amillepora03530-RA | 0.225022424 |
| GO:0043394 | proteoglycan binding | 1 | 1 | 0.01829 | MF | Amillepora03480-RA | 0.225022424 |
| GO:0001594 | trace-amine receptor activity | 1 | 1 | 0.01829 | MF | Amillepora04708-RA | 0.225022424 |
| GO:0047130 | saccharopine dehydrogenase (NADP+, L-lysine-forming) activity | 1 | 1 | 0.01829 | MF | Amillepora16854-RA | 0.225022424 |
| GO:0047131 | saccharopine dehydrogenase (NAD+, L-glutamate-forming) activity | 1 | 1 | 0.01829 | MF | Amillepora16854-RA | 0.225022424 |
| GO:0045545 | syndecan binding | 1 | 1 | 0.01829 | MF | Amillepora03480-RA | 0.225022424 |
| GO:0032557 | pyrimidine ribonucleotide binding | 1 | 1 | 0.01829 | MF | Amillepora22291-RA | 0.225022424 |
| GO:0031151 | histone methyltransferase activity (H3-K79 specific) | 1 | 1 | 0.01829 | MF | Amillepora25070-RA | 0.225022424 |
| GO:0008227 | G protein-coupled amine receptor activity | 1 | 1 | 0.01829 | MF | Amillepora04708-RA | 0.225022424 |
| GO:0032574 | 5'-3' RNA helicase activity | 1 | 1 | 0.01829 | MF | Amillepora18353-RA | 0.225022424 |
| GO:0001565 | phorbol ester receptor activity | 1 | 1 | 0.01829 | MF | Amillepora12560-RA | 0.225022424 |
| GO:0017077 | oxidative phosphorylation uncoupler activity | 1 | 1 | 0.01829 | MF | Amillepora04680-RA | 0.225022424 |
| GO:0003920 | GMP reductase activity | 1 | 1 | 0.01829 | MF | Amillepora10796-RA | 0.225022424 |
| GO:0047690 | aspartyltransferase activity | 1 | 1 | 0.01829 | MF | Amillepora24553-RA | 0.225022424 |
| GO:0120152 | calcium-dependent outer dynein arm binding | 1 | 1 | 0.01829 | MF | Amillepora11351-RA | 0.225022424 |
| GO:0140659 | cytoskeletal motor regulator activity | 1 | 1 | 0.01829 | MF | Amillepora11351-RA | 0.225022424 |
| GO:0097363 | protein O-GlcNAc transferase activity | 1 | 1 | 0.01829 | MF | Amillepora07718-RA | 0.225022424 |
| GO:0008521 | acetyl-CoA transmembrane transporter activity | 1 | 1 | 0.01829 | MF | Amillepora19435-RA | 0.225022424 |
| GO:0002135 | CTP binding | 1 | 1 | 0.01829 | MF | Amillepora22291-RA | 0.225022424 |
| GO:0005545 | 1-phosphatidylinositol binding | 1 | 1 | 0.01829 | MF | Amillepora20340-RA | 0.225022424 |
| GO:0061775 | cohesin loading activity | 1 | 1 | 0.01829 | MF | Amillepora03553-RA | 0.225022424 |
| GO:0004753 | saccharopine dehydrogenase activity | 1 | 1 | 0.01829 | MF | Amillepora16854-RA | 0.225022424 |
| GO:0004832 | valine-tRNA ligase activity | 1 | 1 | 0.01829 | MF | Amillepora01328-RA | 0.225022424 |
| GO:0097729 | 9+2 motile cilium | 16 | 4 | 0.00033 | CC | Amillepora11351-RA, Amil | 0.0357 |
| GO:0016593 | Cdc73/Paf1 complex | 7 | 3 | 0.00034 | CC | Amillepora13525-RA, Amil | 0.0357 |
| GO:0005635 | nuclear envelope | 61 | 7 | 0.00034 | CC | Amillepora23397-RA, Amil | 0.0357 |
| GO:0140534 | endoplasmic reticulum protein-containing complex | 42 | 5 | 0.00208 | CC | Amillepora06449-RA, Amil | 0.13125 |
| GO:0015629 | actin cytoskeleton | 62 | 6 | 0.00223 | CC | Amillepora22084-RA, Amil | 0.13125 |
| GO:0030667 | secretory granule membrane | 13 | 3 | 0.0025 | CC | Amillepora14835-RA, Amil | 0.13125 |
| GO:0031225 | anchored component of membrane | 15 | 3 | 0.00386 | CC | Amillepora09624-RA, Amil | 0.1737 |
| GO:0030135 | coated vesicle | 55 | 5 | 0.00679 | CC | Amillepora15763-RA, Amil | 0.181373684 |
| GO:0009897 | external side of plasma membrane | 7 | 2 | 0.0093 | CC | Amillepora24396-RA, Amil | 0.181373684 |
| GO:0030136 | clathrin-coated vesicle | 21 | 3 | 0.01024 | CC | Amillepora06175-RA, Amil | 0.181373684 |
| GO:0005813 | centrosome | 87 | 6 | 0.01175 | CC | Amillepora22273-RA, Amil | 0.181373684 |
| GO:0005751 | mitochondrial respiratory chain complex IV | 8 | 2 | 0.01222 | CC | Amillepora24987-RA, Amil | 0.181373684 |
| GO:0045277 | respiratory chain complex IV | 9 | 2 | 0.01549 | CC | Amillepora24987-RA, Amil | 0.181373684 |
| GO:0031090 | organelle membrane | 875 | 31 | 0.016 | CC | Amillepora12729-RA, Amil | 0.181373684 |
| GO:0005798 | Golgi-associated vesicle | 25 | 3 | 0.01661 | CC | Amillepora06397-RA, Amil | 0.181373684 |

| | | | | | | | |
|---|---|---|---|---|---|---|---|
| GO:0098796 | membrane protein complex | 307 | 13 | 0.01678 | CC | Amillepora06175-RA, Amil | 0.181373684 |
| GO:0042175 | nuclear outer membrane-endoplasmic reticulum membrane network | 276 | 12 | 0.01763 | CC | Amillepora03625-RA, Amil | 0.181373684 |
| GO:0031965 | nuclear membrane | 26 | 3 | 0.01849 | CC | Amillepora26224-RA, Amil | 0.181373684 |
| GO:0005789 | endoplasmic reticulum membrane | 247 | 11 | 0.01934 | CC | Amillepora03625-RA, Amil | 0.181373684 |
| GO:0098827 | endoplasmic reticulum subcompartment | 250 | 11 | 0.02095 | CC | Amillepora03625-RA, Amil | 0.181373684 |
| GO:0001533 | cornified envelope | 1 | 1 | 0.02188 | CC | Amillepora20112-RA | 0.181373684 |
| GO:0042599 | lamellar body | 1 | 1 | 0.02188 | CC | Amillepora14835-RA | 0.181373684 |
| GO:0070765 | gamma-secretase complex | 1 | 1 | 0.02188 | CC | Amillepora15763-RA | 0.181373684 |
| GO:0098894 | extrinsic component of presynaptic endocytic zone membrane | 1 | 1 | 0.02188 | CC | Amillepora20340-RA | 0.181373684 |
| GO:0099243 | extrinsic component of synaptic membrane | 1 | 1 | 0.02188 | CC | Amillepora20340-RA | 0.181373684 |
| GO:0036501 | UFD1-NPL4 complex | 1 | 1 | 0.02188 | CC | Amillepora22025-RA | 0.181373684 |
| GO:0042589 | zymogen granule membrane | 1 | 1 | 0.02188 | CC | Amillepora15763-RA | 0.181373684 |
| GO:1902560 | GMP reductase complex | 1 | 1 | 0.02188 | CC | Amillepora10796-RA | 0.181373684 |
| GO:0120219 | subapical part of cell | 1 | 1 | 0.02188 | CC | Amillepora00666-RA | 0.181373684 |
| GO:0097227 | sperm annulus | 1 | 1 | 0.02188 | CC | Amillepora22151-RA | 0.181373684 |
| GO:0098888 | extrinsic component of presynaptic membrane | 1 | 1 | 0.02188 | CC | Amillepora20340-RA | 0.181373684 |
| GO:0042824 | MHC class I peptide loading complex | 1 | 1 | 0.02188 | CC | Amillepora00093-RA | 0.181373684 |
| GO:0097232 | lamellar body membrane | 1 | 1 | 0.02188 | CC | Amillepora14835-RA | 0.181373684 |
| GO:0097233 | alveolar lamellar body membrane | 1 | 1 | 0.02188 | CC | Amillepora14835-RA | 0.181373684 |
| GO:0098833 | presynaptic endocytic zone | 1 | 1 | 0.02188 | CC | Amillepora20340-RA | 0.181373684 |
| GO:0098835 | presynaptic endocytic zone membrane | 1 | 1 | 0.02188 | CC | Amillepora20340-RA | 0.181373684 |
| GO:0097208 | alveolar lamellar body | 1 | 1 | 0.02188 | CC | Amillepora14835-RA | 0.181373684 |
| GO:0034098 | VCP-NPL4-UFD1 AAA ATPase complex | 1 | 1 | 0.02188 | CC | Amillepora22025-RA | 0.181373684 |
| GO:0099512 | supramolecular fiber | 160 | 8 | 0.02386 | CC | Amillepora12203-RA, Amil | 0.192715385 |
| GO:0099081 | supramolecular polymer | 163 | 8 | 0.02631 | CC | Amillepora12203-RA, Amil | 0.194727273 |
| GO:0005622 | intracellular anatomical structure | 4413 | 116 | 0.02662 | CC | Amillepora11097-RA, Amil | 0.194727273 |
| GO:0005930 | axoneme | 30 | 3 | 0.0271 | CC | Amillepora22151-RA, Amil | 0.194727273 |
| GO:0030137 | COPI-coated vesicle | 12 | 2 | 0.0272 | CC | Amillepora15763-RA, Amil | 0.194727273 |
| GO:0005884 | actin filament | 12 | 2 | 0.0272 | CC | Amillepora12203-RA, Amil | 0.194727273 |
| GO:0097014 | ciliary plasm | 31 | 3 | 0.02953 | CC | Amillepora22151-RA, Amil | 0.20671 |
| GO:0032154 | cleavage furrow | 14 | 2 | 0.03645 | CC | Amillepora24931-RA, Amil | 0.244292553 |
| GO:0070069 | cytochrome complex | 14 | 2 | 0.03645 | CC | Amillepora24987-RA, Amil | 0.244292553 |
| GO:0005940 | septin ring | 2 | 1 | 0.04329 | CC | Amillepora22151-RA | 0.247933636 |
| GO:0055044 | symplast | 2 | 1 | 0.04329 | CC | Amillepora22668-RA | 0.247933636 |
| GO:0035371 | microtubule plus-end | 2 | 1 | 0.04329 | CC | Amillepora05723-RA | 0.247933636 |
| GO:0009506 | plasmodesma | 2 | 1 | 0.04329 | CC | Amillepora22668-RA | 0.247933636 |
| GO:0042588 | zymogen granule | 2 | 1 | 0.04329 | CC | Amillepora15763-RA | 0.247933636 |
| GO:0140445 | chromosome, telomeric repeat region | 2 | 1 | 0.04329 | CC | Amillepora21960-RA | 0.247933636 |
| GO:0002177 | manchette | 2 | 1 | 0.04329 | CC | Amillepora21990-RA | 0.247933636 |
| GO:1990752 | microtubule end | 2 | 1 | 0.04329 | CC | Amillepora05723-RA | 0.247933636 |
