## Supplementary Material 4 for "Synergistic genomic mechanisms of adaptation to ocean acidification in a coral holobiont"

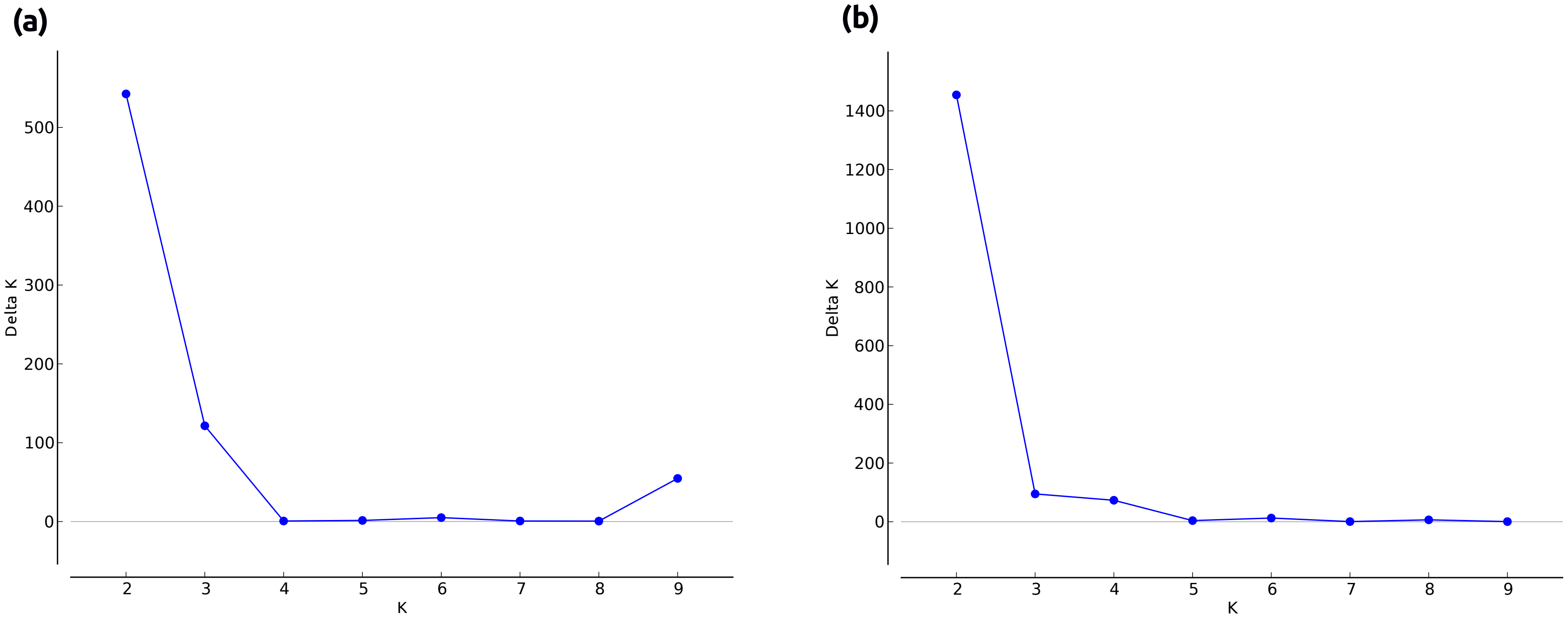


**Supplementary Material 4:** Delta *K* plots for the STRUCTURE analyses, (a) using the 11,169 independent neutral SNP dataset and (b) using the 625 cadidate adaptive SNP dataset.
