## Supplementary Material 5 for "Synergistic genomic mechanisms of adaptation to ocean acidification in a coral holobiont"

**Supplementary material 5:** REVIGO treemaps for Biological Process, Cellular Component, and Molecular Function

Revigo TreeMap Biological Process

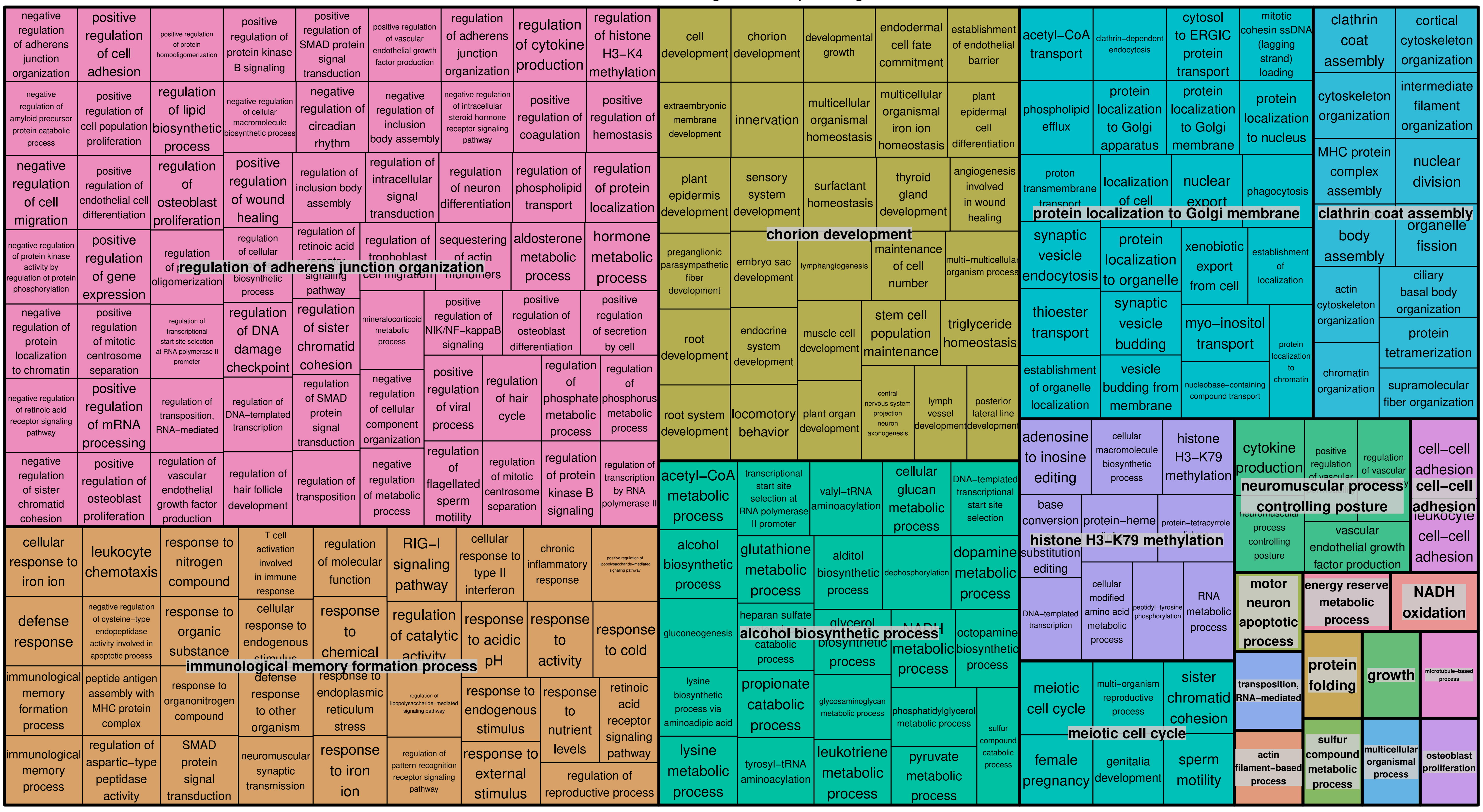


Revigo TreeMap Cellular Component

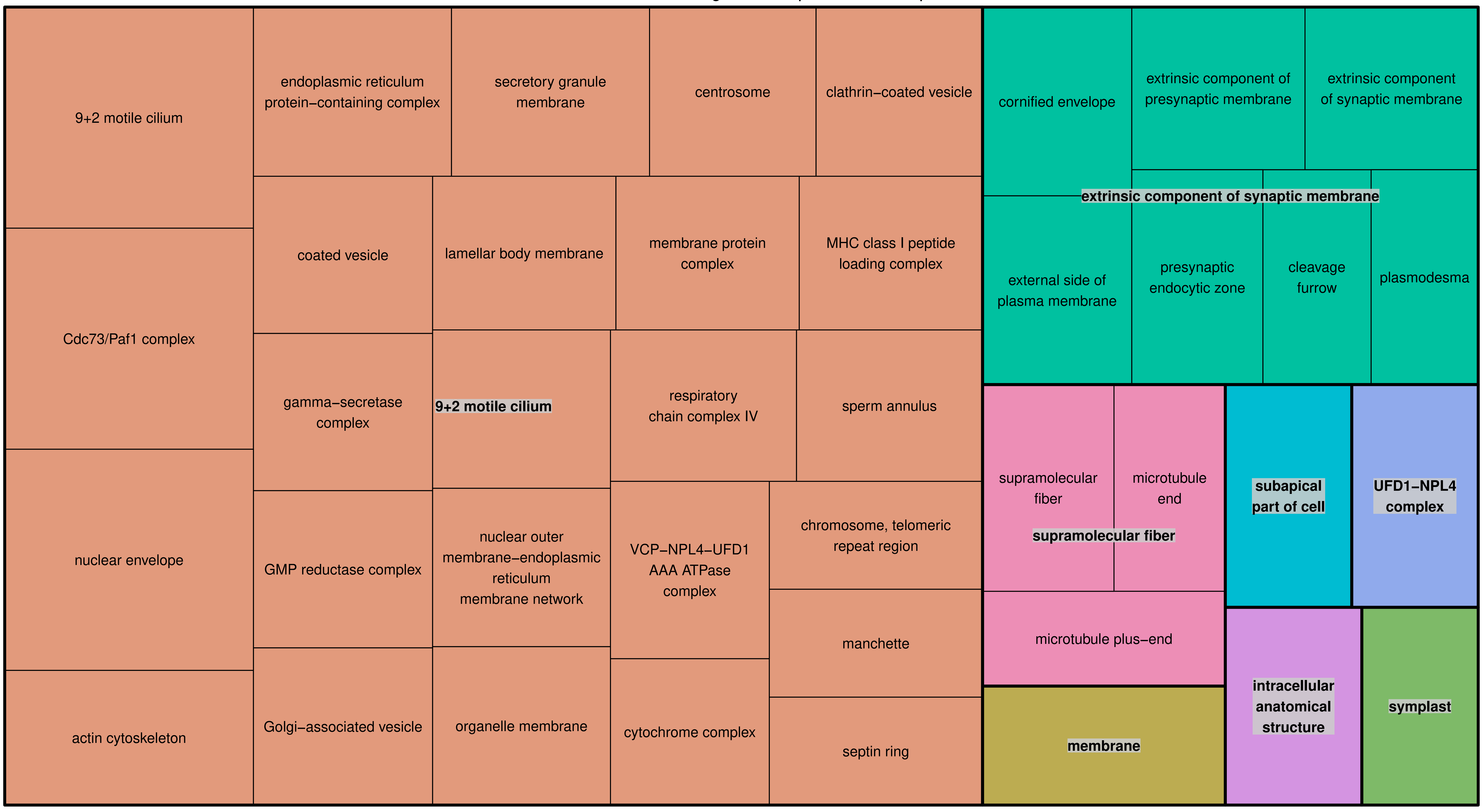


Revigo TreeMap Molecular Function

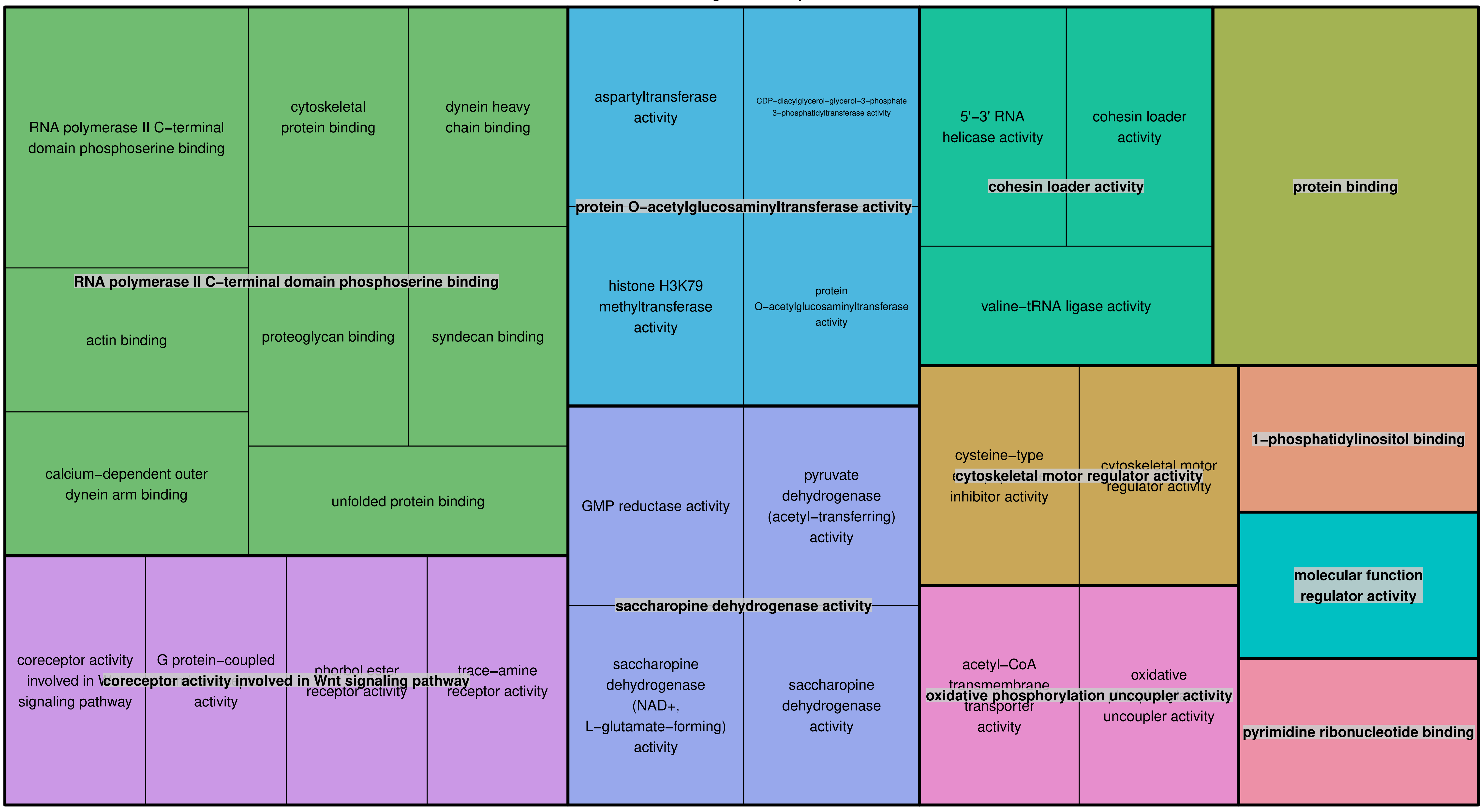
